# Integrating multi-modal transcriptomics identifies cellular subtypes with distinct roles in PDAC progression

**DOI:** 10.1101/2025.02.11.637627

**Authors:** Jun Wu, Tenghui Dai, Ziyue Li, Meng Pan, Wei Zhang, Hao Chen, Guansheng Zheng, Li Qiao, Qizhou Lian, Yang Liu, Jierong Chen

## Abstract

**Background:** Pancreatic ductal adenocarcinoma (PDAC) features a complex tumor microenvironment (TME) that significantly influences patient outcomes. Understanding the TME’s cellular composition and interactions is crucial for identifying therapeutic targets to improve treatment.

**Methods:** We performed an integrative analysis combining 88 single-cell RNA sequencing (scRNA-seq) samples with 187,520 cells, 20 Visium spatial transcriptomics (ST-seq) samples with 67,933 spots, and 1383 bulk RNA-seq samples and 2 Xenium high-resolution ST-seq samples with 307,679 cells, to delineate and characterize distinct subpopulations of fibroblasts, macrophages, T/NK cells, and B/plasma cells within the PDAC microenvironment. Correlations among major cell types across 12 PDAC datasets were assessed through gene set variation analysis (GSVA). Kaplan-Meier survival analysis was used to evaluate the prognostic significance of cell subtypes. Furthermore, multiplex immunohistochemistry (mIHC) and the Xenium platform was utilized to validate cellular interactions in human PDAC tissues at both protein and RNA levels.

**Results:** Six fibroblast subtypes and eight macrophage/monocyte subtypes were identified. POSTN ^high^ fibroblasts and SPP1 ^high^ macrophages highly infiltrated tumor tissues and were associated with poor prognosis. Most immune cell subtypes mediate adverse prognoses, except for CCL4 ^high^ CD8+ T _EFF_ cells and IGHG1 ^high^ plasma cells, which are linked to favorable outcomes. ST-seq revealed spatial colocalization of POSTN ^high^ Fibro and SPP1 ^high^ Macro cells, as well as CCL4 ^high^ CD8+ T _EFF_ and IGHG1 ^high^ plasma cells. These findings were corroborated by mIHC and validated using Xenium spatial transcriptomics with single-cell resolution, which confirmed the expression and spatial proximity of these markers at both the gene and protein levels.

**Conclusion:** This integrative analysis of the PDAC TME underscores the prognostic importance of POSTN ^high^ fibroblasts and SPP1 ^high^ macrophages, while also highlighting the protective roles of CCL4 ^high^ CD8+ T _EFF_ cells and IGHG1 ^high^ plasma cells. These insights could improve PDAC treatment strategies.

## Introduction

Pancreatic ductal adenocarcinoma (PDAC) stands as the most prevalent and aggressive subtype of pancreatic cancer, accounting for approximately 90% of all cases. Originating primarily as tubular adenocarcinomas from the ductal epithelium, PDAC remains a formidable challenge in oncology due to its exceedingly poor prognosis and limited therapeutic options [1, 2]. Projections suggest that by 2030, PDAC will become the second leading cause of cancer-related mortality [2]. The lethality of PDAC is exacerbated by the absence of effective early detection methods, which often results in diagnoses at advanced stages, with a staggering 80% of cases identified at a locally advanced stage accompanied by widespread metastases, particularly to the liver. This leaves the majority of patients ineligible for surgical resection, contributing to a dismal five-year survival rate of approximately 11% [3–6]. As a result, there is an urgent need for the development of early detection strategies and more effective therapeutic interventions for PDAC.

The PDAC tumor microenvironment (TME) is particularly complex, and is characterized by a dense stroma that plays crucial roles in tumor progression, immune evasion, and resistance to therapy [7, 8]. This stromal compartment comprises a variety of cellular elements, including cancer-associated fibroblasts (CAFs), the extracellular matrix (ECM), and a diverse array of immune cells, all of which contribute to the intricate dynamics of PDAC pathobiology [7]. Despite advances in systemic chemotherapy, which remains the mainstay of treatment aimed at prolonging survival, efforts to target the stromal barrier—such as the inhibition of the fibroblast-associated albumin-binding protein SPARC to increase the delivery of gemcitabine—have not yielded the anticipated clinical benefits. Preclinical investigations have failed to substantiate the stromal depletion hypothesis or establish SPARC as a reliable predictive biomarker for treatment efficacy [9–12]. These challenges underscore the necessity for a deeper and more nuanced understanding of the cellular constituents of the TME and their contributions to PDAC progression and therapeutic resistance.

Recent advancements in high-throughput sequencing technologies have markedly expanded our understanding of the cellular heterogeneity and molecular complexity of cancers, including PDAC. Single-cell RNA sequencing (scRNA-seq) [13, 14] and spatial transcriptomics (ST-seq) [15, 16] have emerged as powerful tools that provide unprecedented resolution in dissecting the cellular composition and spatial organization of tumors. These technologies have been instrumental in revealing the intricate cellular interactions and molecular pathways that drive tumorigenesis across various malignancies, including pancreatic [17, 18], lung [19, 20], liver [21–23], breast [24, 25], and prostate [26] cancers. In the context of PDAC, Yousuf *et al*. utilized an integrated scRNA-seq and ST-seq approach to map the immunosuppressive landscape, revealing distinct subsets of cytotoxic T cells that are functionally impaired within the tumor microenvironment [18]. The interplay between immune cells and other tumor-associated cellular components, such as CAFs and tumor-associated macrophages (TAMs), plays a pivotal role in shaping the TME and influencing therapeutic outcomes [27, 28]. For example, CAFs have been shown to modulate the immune landscape by secreting chemokines that inhibit the infiltration of effector T cells, such as CD8+ cytotoxic T lymphocytes, into the tumor core [29]. TAMs, on the other hand, are implicated in promoting tumor proliferation, metastasis, and angiogenesis, thereby reinforcing the immunosuppressive and pro-tumorigenic milieu of PDAC [30]. The synergistic activities of CAFs and TAMs are thought to drive tumor progression through the activation of oncogenic signaling pathways, such as the Akt and STAT pathways, which have been implicated in enhancing the invasive potential of various cancers, including neuroblastoma [31]. Furthermore, Zhang *et al*. demonstrated that CAFs can promote PDAC progression by secreting macrophage colony-stimulating factor, which skews macrophages toward an immunosuppressive M2 phenotype [32]. Despite these advances, the precise roles and therapeutic potential of distinct CAF and TAM subpopulations in the PDAC microenvironment remain inadequately characterized, necessitating further investigation.

In this study, we aimed to delineate the cellular landscape of the PDAC microenvironment, with a particular focus on identifying key cellular subpopulations and potential therapeutic targets. By integrating scRNA-seq, ST-seq, and bulk RNA-seq data from diverse cohorts, we comprehensively profiled the infiltration patterns and functional properties of CAFs, TAMs, and immune cell subpopulations within the PDAC TME. We also correlated these findings with patient prognosis and validated key cellular interactions via multiplex immunohistochemistry (mIHC) and Xenium spatial transcriptomics with single-cell resolution. Our analysis aimed to identify novel prognostic markers and therapeutic targets while providing insights into the immunosuppressive mechanisms at play within the PDAC microenvironment. Through the integration of publicly available datasets, this study aims to contribute to the development of more effective therapeutic strategies for the management of PDAC.

## Methods

### Data collection

88 samples of scRNA-seq data were obtained from five research cohorts, including PDAC, liver metastases (LM), PDAC adjacent-tumor (AT), and normal pancreas (NP; Supplementary Table S1) cohorts. 20 samples of ST-seq data were derived from three research cohorts (Supplementary Table S2), and 1383 samples of bulk RNA-seq gene expression data were collected from twelve research cohorts (Supplementary Table S3). Among these, the TCGA-PAAD and ICGC cohorts were also used to evaluate the associations of genes or cellular subtypes with patient prognosis.

The Xenium spatial transcriptomics, a high-resolution spatial genomics technique with single-cell resolution, was employed in this study. Two samples, selected to meet the objectives of this research, were collected. One sample was derived from the FFPE Human PDAC dataset available in the 10x Genomics database, specifically the Human Immuno-Oncology Profiling Panel (https://www.10xgenomics.com/datasets/ffpe-human-ductal-adenocarcinoma-data-with-human-immuno-oncology-profiling-panel-1-standard). The second sample was obtained from the GEO database, consisting of a PDAC surgical specimen associated with high-grade pancreatic intraepithelial neoplasia (PanIN) (GSE267680, GSM8273021)[33].

### ScRNA-seq data processing and integration

The gene expression matrices and cell barcode information were downloaded from databases. The scRNA-seq data included GSE154778, GSE155698, GSE197177, and GSE212966 from the GEO database, and CRA001160 from the GSA database. The analysis was conducted via the R package Seurat (v4.3.0 in R4.1.1, https://satijalab.org/seurat) [34]. The first step involved filtering out low-quality cells with a cutoff value of less than 200 total feature RNA, fewer than 1000 total RNA counts, and more than 10% mitochondrial RNA. The gene expression matrices for each cell were subsequently normalized via SCTransform. Batch correction was performed via the Harmony package to obtain a unified PDAC scRNA-seq project [35].

For the PDAC scRNA-seq project, principal component analysis (PCA) was utilized to reduce dimensionality, followed by uniform manifold approximation and projection (UMAP). 22 clusters were identified using the FindNeighbors (dims=1:30) and FindClusters (resolution=0.3) functions. The average gene expression of well-known markers, including immune cells (*PTPRC*), epithelial cells (*EPCAM*), T/NK (natural killer) cells (*CD3D*, *CD3E*, and *GNLY*), B/plasma cells (*CD79A*, *MS4A1*, and *IGHA1*), mast cells (*CPA3*), macrophages/monocytes (*CD68* and *CD14*), dendritic cells (*CD1C*), fibroblasts (*COL1A1* and *DCN*), endothelial cells (*PECAM1* and *VWF*), and pericytes (*ACTA2* and *CSPG4*), was subsequently evaluated. We annotated a total of 18,7520 cells across 88 samples with 9 major cell types after removing doublets.

### Processing of spatial transcriptomics data

Spatial transcriptomics (ST-seq) data, including the HTAN-PDAC and GEO datasets GSE233293 and GSE202740, totaling 20 ST-seq samples, were jointly analyzed via the R packages Seurat and BayesSpace in R4.1.1 [36]. The gene expression matrices per spot were processed using the standard workflow of BayesSpace, including functions such as spatialPreprocess, spatialCluster, and clusterPlot. Specific parameter settings are provided upon request. The spatial cluster information obtained from BayesSpace was integrated into Seurat for subsequent analysis.

Spatial expression patterns of cell subtypes are presented by calculating the expression of representative genes and well-known markers. Image enhancement visualization is performed using embedded functions in BayesSpace (https://github.com/edward130603/BayesSpace).

### Differential gene analysis

For the differential expressed genes (DEGs) analysis of each cell type or cell subtype in the scRNA-seq data, the FindAllMarkers function in Seurat was utilized, with the parameters logfc.threshold=0.25, only.pos=T, and test.use=wilcox. Default values were used for other parameters. The R package pheatmap was employed for visualizing gene expression heatmaps after scaling.

For bulk RNA-seq data, differential expression analysis of genes was performed using the R package DESeq2, considering appropriate grouping conditions. Gene expression values were treated as count data for DEG calculation, which was then utilized for subsequent downstream analysis.

### Functional enrichment analysis

For the functional enrichment analysis of the eight cell types in the tumor microenvironment (TME) using Hallmark gene sets obtained from the Molecular Signatures Database (MSigDB, https://www.gsea-msigdb.org/gsea/msigdb), the following steps were performed: 1) The GSVA software package was employed to compute the single-sample gene set enrichment analysis (ssGSEA) scores for 50 hallmark gene sets per cell. 2) Differential expression analysis was conducted between cancer (PDAC primary + liver metastases) and normal (normal pancreas + PDAC adjacent) cells at the scRNA level. 3) significance of differential expression per hallmark was determined using Student’s t test. 4) Visualization of the results was carried out via the ggplot2 package.

Similarly, for the subgroups of fibroblasts and macrophages/monocytes, enrichment analysis of 14 cancer hallmark gene sets was conducted via the R package GSVA. The scaled enrichment scores were visualized using the R package pheatmap.

For bulk RNA-seq, gene set enrichment analysis (GSEA) of hallmark and gene ontology biological process (GO BP) terms was performed using the standard workflow of the R package fgsea. TCGA-PAAD cancer samples were grouped on the basis of the combination of POSTN and SPP1, or CCL4 and ICHG1, and ranked according to the fold change in gene expression. The normalized enrichment scores (NESs) were visualized using the ggplot2 package in R4.1.1.

### Cell-cell correlation analysis

The cell-cell correlation analysis between different cell types is based on multiple datasets. First, the top expressed genes for each cell type or cell subgroup were identified through differential analysis. The top 10-20 genes were subsequently selected as representatives for each cell type (Supplementary Table S4). Then, GSVA scores for each cell type were computed for each sample across the 12 bulk RNA-seq datasets. Finally, Spearman correlation coefficients between cells are calculated, and the results are visualized using ggplot2 with multiple styles. The spatial correlation is calculated as the Spearman coefficient of gene expression for each spot, even though a spot may contain multiple or different cell types.

### Survival analysis

Survival analysis was conducted on the basis of clinical information from PDAC patients from the TCGA and ICGC databases. For single-gene survival analysis, the R package survminer was used to calculate the optimal cutoff threshold, dividing patients into two groups. For dual-gene survival analysis, optimal cutoff values for each gene are determined first, and then samples are divided into two groups based on high and low expression levels of both genes. For survival analysis of cell types or subtypes, GSVA enrichment scores for the cell type per tumor sample were initially calculated, followed by partitioning into two groups based on the optimal cutoff threshold of scores. The significance of survival analysis was assessed using log-rank tests. Kaplan-Meier plots were visualized via the R package ggsurvplot function.

### Transcription factor regulation analysis

The activation of transcription factor regulons was computed using pySCENIC (https://github.com/aertslab/pySCENIC). The gene expression matrices and cell type information for fibroblasts and macro/monocytes were extracted from Seurat and passed to pySCENIC. The transcription factor activity is subsequently predicted for fibroblast and macro/monocyte subtypes using a standard pipeline. The results of the regulons can be found in supplementary Tables S5 and S6. The relative expression of the regulons was then visualized using R package pheatmap.

### Multiplexed immunohistochemistry (mIHC) staining

Paraffin-embedded tissue blocks from patients with pathologically confirmed pancreatic ductal adenocarcinoma were collected from the Second Affiliated Hospital of Wannan Medical College. The tissues were divided into two groups. After deparaffinization and dehydration, the tissues were repaired with a repair solution (citric acid pH 6.0 / EDTA pH 9.0) at high temperature and pressure for 2 minutes. The mixture was allowed to cool naturally, and then washed with TBS three times for 5 minutes each. The samples were incubated with 10% donkey serum (antgene ANT050) at room temperature for 30 minutes. In one group, the following primary antibodies were added: POSTN1 (PA5-34641, Invitrogen, 1:1000), a-SMA (ab124964, Abcam, 1:1000), CK19 (ab124864, Abcam, 1:1000), osteopontin (PA5-34579, Invitrogen, 1:1000), CD163 (ab182422, Abcam, 1:1000). In the other group, the corresponding primary antibodies were added: CD3 (ab16669, Abcam, 1:2000), CD8 (ab237709, Abcam, 1:1000), CCL4 (PA5-34509, Invitrogen, 1:500), CD38 (PA5-82791, Invitrogen, 1:500), and IGHG1 (PA5-75428, Invitrogen, 1:1000). The mixture was placed in a refrigerator at 4 °C overnight. The next day, the slides were removed and heated to room temperature for 15 minutes. The slides were placed in PBS and washed 3 times for 5 minutes each. After drying, an IgG H&L (HRP; ab205718) secondary antibody conjugated to the corresponding species of the primary antibody was added to the circles on the slides, which were subsequently incubated at room temperature for 50 minutes. The slides were washed again in PBS 3 times for 5 minutes each. Tyramide signal amplification (TSA) working solution was added according to the TSA kit instructions, and the slides were incubated at room temperature in the dark for 50 minutes. In the presence of HRP, the tyramide fluorophore is activated and covalently binds to tyrosine residues of the target protein, amplifying the signal. The noncovalently bound antibodies were removed via microwave treatment, and the above steps were repeated with different TSA-derived fluorescent dyes for multicolor staining. The TSA-derived fluorescent dyes added sequentially in each step are CY3-tyramide (G1223, Servicebio, 1:500), iF594-tyramide (G1242, Servicebio, 1:500), FITC-tyramide (G1222, Servicebio, 1:500), iF700-Tyramide (G1232, Servicebio, 1:500), and iF440-Tyramide (G1250, Servicebio, 1:500). The samples were washed with TBS three times for 5 minutes each. After drying, DAPI staining solution was added, and the samples were incubated in the dark at room temperature for 10 minutes. After drying, the slides were treated with an autofluorescence quencher for 5 minutes, and rinsed with running water for 10 minutes. The samples were washed with TBS three times for 5 minutes each, dried, and mounted with anti-fade mounting medium. Images were captured via a multispectral imaging system (Pannoramic MIDI, 3DHISTECN). On the basis of fluorescence intensity and distribution, qualitative, locational, and semiquantitative analyses of the target proteins can be performed.

### Analysis of High-dimensional spatial transcriptomics (Xenium)

For the two samples analyzed using the Xenium platform with single-cell resolution RNA in situ hybridization, Seurat V5 was employed for the analysis with identical parameters. The processed data was downloaded and used as input for Seurat. The analysis pipeline began with the removal of cells with zero counts. Next, the data was normalized using SCTransform. After performing PCA, FindNeighbors and FindClusters (resolution = 0.1) were applied to classify spatial cell clusters. The ImageDimPlot function was used to visualize the spatial distribution of regions and modules. Detailed parameters can be found in the Seurat V5 official tutorial for analyzing Xenium sequencing data.

For the spatial region identification of the PDAC surgical specimen with associated high-grade PanIN sample with 72,822 cells, immune-related genes and stroma-related genes were utilized, alongside markers identified by the original authors for high-grade PanIN, to ensure accurate region classification (refer to Supplementary Figure S10 for details). For the fibroblast clusters, cells were classified into POSTN ^high^ clusters when POSTN expression exceeded the mean POSTN expression level. For the FFPE Human PDAC with Human Immuno-Oncology Profiling Panel sample with 234,857 cells, spatial region identification was performed using immune-related genes, stroma-related genes, and tumor-associated genes (refer to Supplementary Figure S10 for details).

### Statistical analysis

When appropriate, intergroup differences were assessed via either Student’s t test or the Wilcoxon test, depending on the data distribution. Survival analysis was conducted using the log-rank test. Regardless of the test used, a p value less than 0.05 is considered significant and denoted by *, whereas a p value greater than 0.05 is considered not significant and denoted by “ns”. When p < 0.01, it is represented as **, p < 0.001 as ***, and p < 0.0001 or lower as ****.

## Results

### A single-cell and spatial transcriptomics atlas of human PDAC tissues

To decipher the cellular composition of the PDAC tumor microenvironment, scRNA-seq data were collected from five independent cohorts, including PDAC primary tumor (n=60), liver metastases (LM; n=10), PDAC adjacent-tumor (AT; n=6), and normal pancreas (NP; n=12) samples. Additionally, spatial transcriptomic samples from PDAC patients (n=20) were collected (Fig. 1A). After the scRNA-seq data were filtered to remove low-quality and suspected doublet cells, a total of 187,520 cells from 88 samples were retained for further analysis (Fig. 1B and S1). After normalizing gene expression, we applied principal component analysis (PCA) based on highly variable genes. To correct for batch effects in the scRNA-seq samples, we further applied Harmony-corrected PCA to generate a UMAP embedding space (Fig. 1B-D). we subsequently performed graph-based clustering and annotated each cluster with their representative markers (Fig. S1C). The cells were classified into 9 major cell types (Fig. 1D and S1D), including epithelial cells identified by the expression of EPCAM; T/NK cells marked by CD3D and CD3E; B/plasma cells expressing MS4A1, CD79A, and IGHA1; macro/mono (macrophage/monocyte) cells identified by CD68 and CD14; dendritic cells (DCs) marked by CD1C; mast cells marked by CPA3; fibroblasts positive for COL1A1 and DCN; endothelial cells marked by PECAM and VWF; and pericytes marked by ACTA2 and CSPG4. Simultaneously, spatial spots were effectively categorized into distinct clusters via BayesSpace, facilitating the mapping of the spatial abundance of features and gene expression (Fig. S2). All major cell types exhibited variable cell fractions across tissue sources, revealing heterogeneity in the cellular composition of the NP, AT, PDAC, and LM (Fig. 1E). Across all the samples, the proportions of DC cells and mast cells were relatively low. We observed that the proportion of infiltrating fibroblasts at LM sites was significantly lower than that in PDAC primary tumors. The infiltration proportion of macrophages in PDAC primary tumors did not significantly differ from that in LM or AT but was higher than that in NP. The infiltration proportions of B cells and T/NK cells in PDAC primary tumors were greater than those in NP sites. Our analysis highlights the distinct cellular heterogeneity within the PDAC tumor microenvironment, revealing key differences between primary tumors, metastatic lesions, and PDAC adjacent tissues that may offer insights into PDAC progression and therapeutic response.

**Figure 1.**
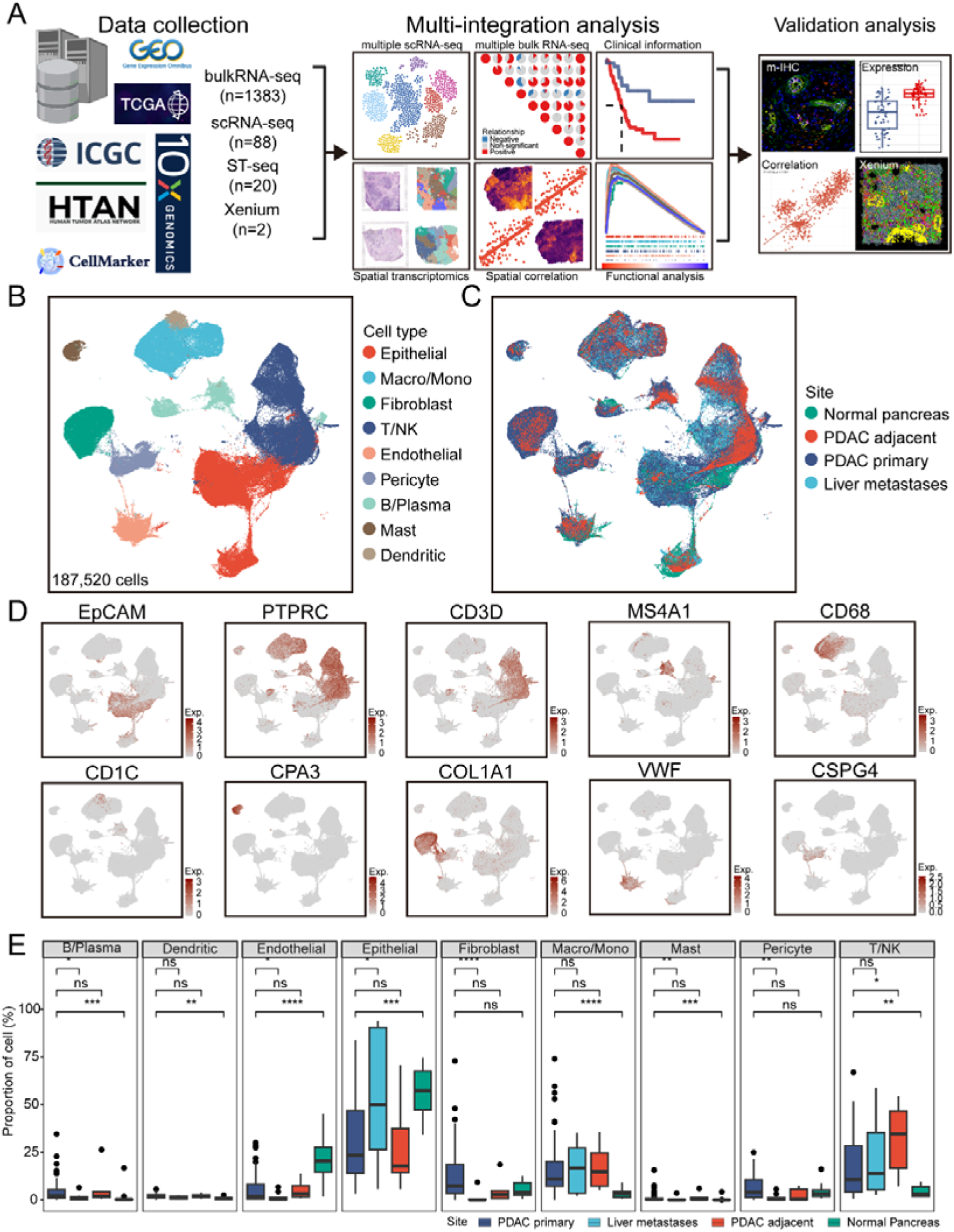
Workflow and overview of a multi-data atlas for human PDAC. (A). Graphic overview of the study design. Single-cell RNA sequencing (scRNA-seq) data related to pancreatic ductal adenocarcinoma (PDAC), along with spatial transcriptomics (ST), bulk RNA sequencing (BulkRNA-seq), and clinical information, were obtained from the GEO, GSA, TCGA, ICGC, HTAN and 10X Genomics databases for bioinformatics analysis. The identified biological markers and targets were further validated via multiplex immunofluorescence staining and Xenium high-resolution spatial transcriptomics. (B) UMAP plots of 187,520 cells from 88 PDAC patient samples, showing nine major cell clusters. Each cluster is represented in a different color. (C) UMAP plots of samples collected from normal pancreas, adjacent tissues, liver metastasis, and primary PDAC sites. Each site is depicted in a distinct color. (D) UMAP plots showing the expression levels of selected known marker genes. (E) Bar plots displaying the proportions of nine cell types across the four different sites. Statistics based on the Wilcoxon test. p<0.05 was considered a statistically significant difference, p<0.01**, p<0.001***, p<0.0001 ****.

### Characteristics of the cellular composition and intercellular interactions within the PDAC microenvironment and prediction of clinical outcome

To decipher the regulation of infiltrating cell subsets in PDAC, we leveraged hallmark gene sets from the MsigDB to analyze pathway alterations between tumor (PDAC primary and LM) and PDAC adjacent normal (NP and AT) tissues across fibroblast, macrophages/monocytes, T/NK, endothelial, pericyte, B/plasma, mast, and dendritic cell populations (Fig. 2A). Compared with that in normal tissues, the epithelial-mesenchymal transition (EMT) pathway is enriched in endothelial, pericyte, fibroblast and macrophages/monocytes populations in tumors, indicating the involvement of endothelial cells and fibroblasts in the progression of EMT in PDAC (Fig. 2A). The immune-related pathways, including IL2/STAT5 signaling, IL6/JAK/STAT3 signaling, and the inflammatory response, were enriched not only in immune cells such as T/NK cells and macrophages/monocytes but also in pericytes, endothelial cells, and fibroblasts from tumors compared to normal samples (Fig. 2A). These findings suggest that these cells might participate in the immune processes within the PDAC microenvironment. Notably, cell cycle regulation-related pathways, including E2F targets, the G2M checkpoint, and the P53 pathway, were significantly enriched in T/NK cells, pericytes, fibroblasts, and macrophages/monocytes in tumors compared with those in normal tissue (Fig. 2A). Notably, the hypoxia signaling in the PDAC microenvironment was enriched in macrophage/monocyte, which may reflect the involvement of macrophages in reshaping the PDAC microenvironment within hypoxic regions of the tumor (Fig. 2A). We also compared hallmark pathways between PDAC primary and PDAC adjacent normal (NP and AT) tissues, and the pathway changes observed were similar to those between tumor (PDAC primary and LM) and PDAC adjacent normal (NP and AT) tissues (Fig. S3A). Collectively, these findings suggest that the regulatory pathways of major cell types are influenced within the PDAC microenvironment.

**Figure 2.**
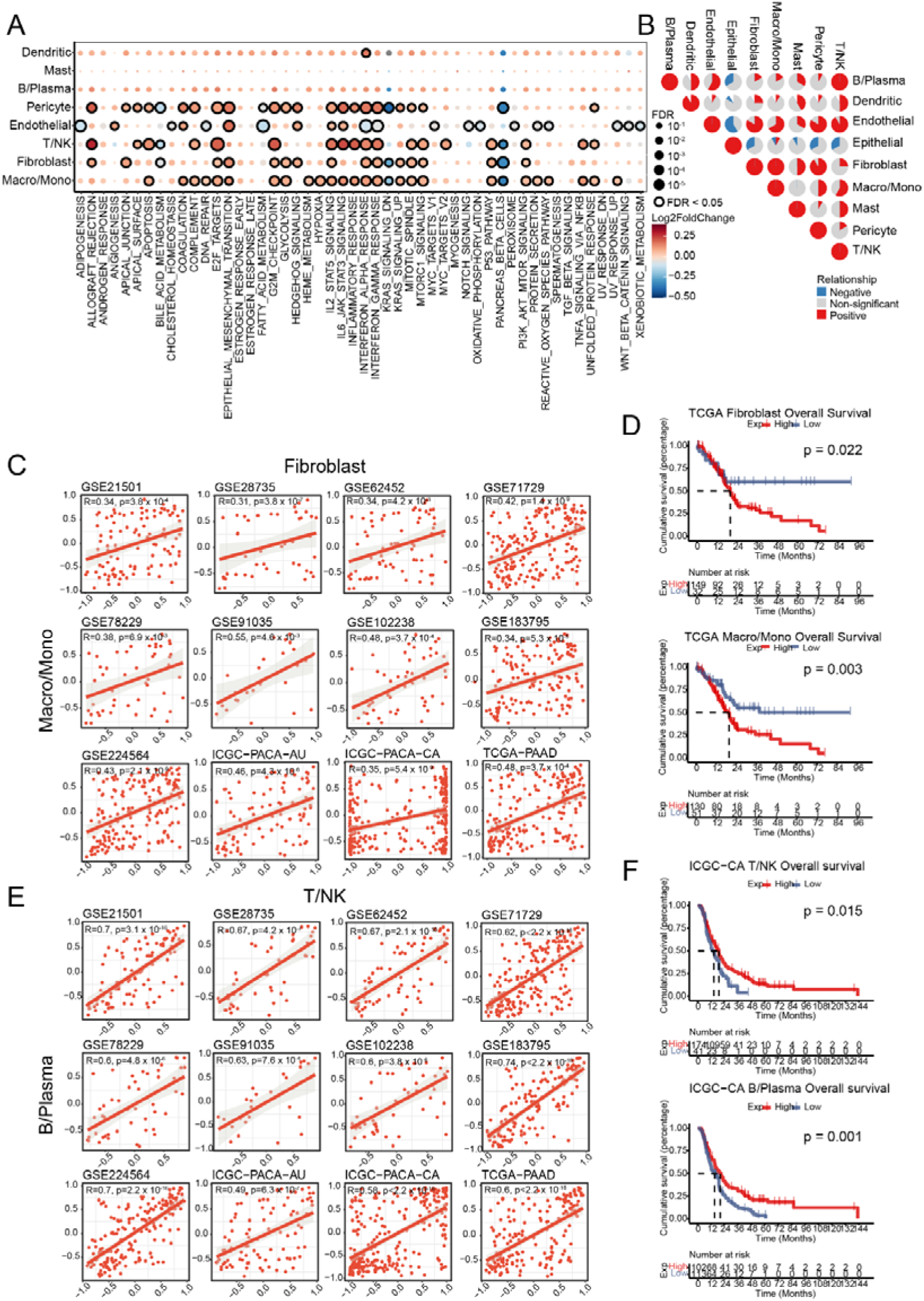
Different cell type infiltration and its association with distinct prognostic outcomes in PDAC. (A). Dot plots showing 50 hallmarks in different cell types between tumor (PDAC primary and LM) and PDAC adjacent/normal tissues. The intensity represents the average fold change of the hallmark GSVA score in tumor versus adjacent/normal tissues. The dot size indicates the FDR for each hallmark. Student’s t test was used to assess significant differences. (B). The proportion of PDAC cohorts with positive (red), negative (blue), or non-significant (gray) correlations for the infiltration of pairwise cell types across 12 independent PDAC cohorts. Spearman’s correlation was used to calculate the correlation coefficient. A p value < 0.05 and coefficient [R] > 0.3 indicate a positive correlation, whereas a p value < 0.05 and coefficient [R] < −0.3 indicate a negative correlation. Other cases are considered non-significant. (C). Scatter plots show the correlation between the infiltration of fibroblasts and macrophages/monocytes across 12 independent PDAC datasets, including TCGA-PAAD (n=181), GSE21501 (n=102), GSE28735 (n=45), GSE62452 (n=69), GSE71729 (n=191), GSE78229 (n=50), GSE91035 (n=25), GSE102238 (n=50), GSE183795 (n=139), GSE224564 (n=175), ICGC-PACA-AU (n=92), and ICGC-PACA-CA (n=264). The error band represents the 95% confidence interval. (D). The Kaplan-Meier curves showed that patients with higher infiltration of fibroblasts (top) and macrophages/monocytes (bottom) are associated with worse outcomes. (E). Scatter plots illustrate the correlation between the infiltration of T/NK and B/Plasma cells across 12 independent PDAC datasets, including TCGA-PAAD (n=181), GSE21501 (n=102), GSE28735 (n=45), GSE62452 (n=69), GSE71729 (n=191), GSE78229 (n=50), GSE91035 (n=25), GSE102238 (n=50), GSE183795 (n=139), GSE224564 (n=175), ICGC-PACA-AU (n=92), and ICGC-PACA-CA (n=264). The error band represents the 95% confidence interval. (F). Kaplan-Meier curves showed that patients with higher infiltration of T/NK cells (top) and B/Plasma cells (bottom) are associated with better outcomes. A two-sided log-rank test with p<0.05 was considered statistically significant in (D, F).

Furthermore, we constructed gene set files using cell type-specific markers to predict GSVA scores for each cell type in large-scale datasets, including 12 independent cohorts (listed in Table S2) such as TCGA-PAAD. To investigate the relationships among different cell populations within the PDAC microenvironment, we analyzed pairwise Spearman correlations within the GSVA scores of the 8 major cell types across the 12 independent PDAC cohorts. We identified two significant positive correlations across all cohorts, including correlations between fibroblasts and macro/mono, and between T/NK and B/plasma cells (spearman correlation coefficient [|R|] > 0.3 and p-value [p] < 0.05 were considered significant correlations, whereas R > 0.3 and p<0.05 indicated positive correlations, Fig. 2B). The positive correlation between fibroblasts and macro/mono cells ranged from 0.31 in the GSE28735 dataset with 45 samples to 0.55 in the GSE91035 dataset with 25 samples across all 12 cohorts (Fig. 2C). To assess the clinical relevance of infiltrating cell types in the PDAC microenvironment with respect to the overall survival (OS) of patients, we analyzed the correlation between cell type scores and the OS of PDAC patients. These analyses revealed that higher infiltration scores of fibroblasts were associated with poorer OS in the TCGA cohort (log-rank test, p=0.022) and the ICGC cohort (log-rank test, p=0.072; Fig. 2D and S3B). Moreover, patients with higher Macro/mono scores were associated with poorer OS in the TCGA cohort (log-rank test, p=0.003) and the ICGC cohort (log-rank test, p=0.014; Fig. 2D and S3C). Furthermore, a positive correlation between T/NK and B/Plasma was observed across the 12 cohorts, with correlation coefficients ranging from 0.6 in the ICGC-PACA-AU cohort with 92 samples to 0.74 in the GSE183795 dataset with 139 samples (Fig. 2E). High infiltration scores of T/NK cells (log-rank test, p=0.015) and high B/Plasma cell scores (log-rank test, p=0.001) in tumor tissues were associated with better OS of patients in the ICGC cohort (Fig. 2F), although these scores were not significantly correlated in the TCGA cohort (Fig. S3D and S3E). These findings may suggest that fibroblasts and macro/mono cells, along with T/NK and B/Plasma cells, play crucial cooperative roles in orchestrating the reshaping of the PDAC microenvironment, thereby mediating opposing outcomes. These observations suggest that fibroblasts and macro/mono cells, along with T/NK and B/plasma cells, may play pivotal cooperative roles in orchestrating the remodeling of the PDAC microenvironment, thereby mediating distinct patient outcomes.

### Tumor-specific POSTN ^high^ fibroblasts and SPP1 ^high^ macrophages are associated with PDAC progression

The observed differences in cell infiltration between tumor tissue and adjacent normal tissue highlight the pivotal role of the tumor microenvironment in PDAC progression. Fibroblasts and macrophages have been suggested as key stromal cells involved in regulating tumorigenesis and cancer progression. However, the heterogeneity of these cell populations remains challenging to elucidate. Here, we aimed to identify and classify fibroblasts and macrophages within the PDAC microenvironment. First, fibroblasts and macro/mono cells were subsetted separately. Then, dimensionality reduction clustering was performed, followed by appropriate clustering of the cell clusters. For fibroblasts, differential analysis was conducted on the six identified clusters (Fig. S3F). Based on the highly expressed genes in each cluster and specific cellular signature markers previously reported, we identified subtypes of fibroblasts (Fig. S3H). Myofibroblasts were grouped on the basis of high expression of ACTA2, MMP11, MYL9, HOPX, and TPM1. Inflammatory fibroblasts were clustered based on high expression of IL6, PDGFRA, CXCL12, CFD, and CXCL1. Additionally, antigen-presenting fibroblasts exhibited high expression of HLA-B and CD74. We identified inflammatory fibroblasts such as APOD ^high^ Fibro and TFPI2 ^high^ Fibro, myofibroblasts such as POSTN ^high^ Fibro and CST1 ^high^ Fibro, and antigen-presenting fibroblasts such as PGK1 ^high^ Fibro and CCL19^high^ Fibro. POSTN ^high^ Fibro showed high expression of POSTN, FN1, MMP11, and collagen genes such as COL1A1, COL1A2 and COL3A1, which are associated with fibroblast activation and extracellular matrix (ECM) remodeling (Fig. S3F). Next, the macrophage/monocyte were clustered into eight clusters via dimensionality reduction. Differential analysis was performed to calculate the representative highly expressed genes in each cluster (Fig. S3G). When the known macrophage and monocyte markers CD68, CD163, CD14, and FCN1 were combined, the cell clusters were defined (Fig. S3I). The macrophages included HMOX1 ^high^ Macro, G0S2 ^high^ Macro, SPP1 ^high^ Macro, SEPP1 ^high^ Macro, APOC1 ^high^ Macro, and CCL3 ^high^ Macro, whereas the monocytes included the FCN1 ^high^ Mono and S100A12 ^high^ Mono subtypes. SPP1 ^high^ Macro highly expressed SPP1, FN1, CCL2, the matrix metalloproteinase genes MMP9, and MMP12 (Fig. S3G).

Mapping cells to UMAP and analyzing the proportions of cells in tissues, we observed that POSTN ^high^ Fibro cells infiltrated more tumor tissues (PDAC primary and LM) than adjacent normal tissues (NP and PDAC adjacent; Fig. 3A and 3B). POSTN ^high^ Fibro infiltration was significant highest in the primary site of PDAC (P<0.05) and LM (P<0.05), whereas fibroblasts were less abundant in normal and PDAC adjacent sites (Fig. S5A). Moreover, the degree of infiltration of SPP1 ^high^ Macro was greater in tumor tissues than in NP, and AT tissues, whereas LM exhibited macrophage infiltration (Fig. 3C and 3D). To explore the associations between different subsets of fibroblasts and macrophage infiltrations, we designed gene sets to calculate GSVA scores for each subset of cells across 12 independent PDAC cohorts (Table S4). Most fibroblasts were positively correlated with macrophages, which is consistent with previous reports, except for the HMOX1 ^high^ Macro subtype (Fig. S5B). Notably, POSTN ^high^ Fibro was significantly correlated with SPP1 ^high^ macrophages in 11 cohorts (R > 0.3, p < 0.05) and weakly correlated in the GSE21501 cohort (R = 0.25, p < 0.05), with R ranging from 0.25 in the GSE21501 dataset to 0.66 in the ICGC-PACA-CA dataset (Fig. 3E and S5B). We subsequently associated the cell type score with the overall survival (OS) of patients, and found that most fibroblast subtypes were associated with poor prognosis in PDAC patients (Fig. S4). For example, CST1 ^high^ Fibro was elevated in PDAC primary tissues, with significant increase observed in PDAC (p < 0.05), whereas no significant elevation was found in liver metastases (Fig. S5A). This proportion is associated with poorer overall survival in PDAC patients within TCGA (p =0.007) and ICGC-CA (p = 0.024) cohort (Fig 3D and S4). This suggests that CST1 may be a potential prognostic indicator for PDAC patients. Notably, both favorable and unfavorable associations with macrophage subtypes, such as CCL3 Macro (log-rank test, p = 0.002) and SEPP1 Macro (log-rank test, p = 0.021) associated with worse OS in the TCGA cohort (Fig. S4A). However, levels of CCL3 ^high^ Macro (log-rank test, p < 0.001) and SEPP1 ^high^ Macro (log-rank test, p < 0.001) were associated with better OS in the ICGC cohort (Fig. S4B). In particular, POSTN ^high^ Fibro (log-rank test, p = 0.005; Fig. 3F) and SPP1 ^high^ Macro (log-rank test, p = 0.0062; Fig. 3G) were associated with worse OS in the TCGA cohort, and POSTN ^high^ Fibro (log-rank test, p = 0.048; Fig. S5C) and SPP1^high^ Macro (log-rank test, p = 0.066; Fig. S5D) were associated with worse OS in the ICGC cohort. Notably, the results from the ST-seq cohort demonstrated colocalization of POSTN ^high^ fibroblasts and SPP1 ^high^ macrophages in spatial terms with significant spatial correlation (R) in 8 samples (R=0.1, p<0.001 in PDAC20, to R=0.34, p<0.001 in PDAC16), with the highest spatial correlation of 0.34 observed in the PDAC16 sample (Fig. 3H and S5E).

**Figure 3.**
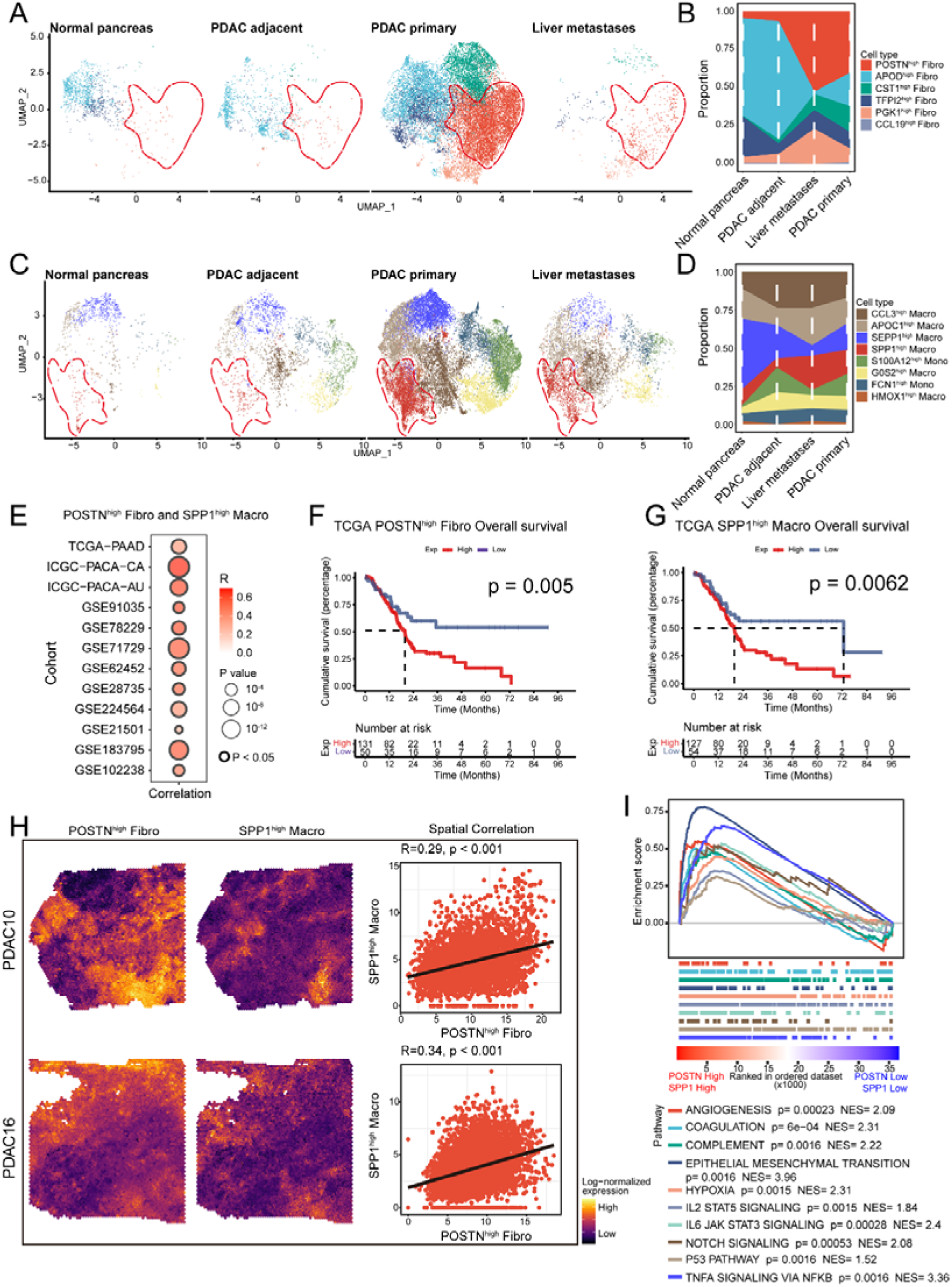
Characterization of POSTN ^high^ fibroblasts and SPP1 ^high^ macrophages and their associations in PDAC TME. (A). UMAP plots displaying the composition of fibroblasts colored by sub cluster. Red dashed circles highlight POSTN ^high^ fibroblasts. (B). The percentage of each fibroblast subclusters across different tissue sites in scRNA-seq. (C). UMAP plots displaying the composition of macrophages/monocytes, colored by subtype. Red dashed circles highlight SPP1 ^high^ macrophages. (D). The percentage of each Macro/Mono subclusters in different tissue sites in scRNA-seq. (E). A dot plot illustrates the Spearman correlation coefficients (R) and their corresponding p-values between POSTN ^high^ fibroblasts and SPP1 ^high^ macrophages across 12 independent cohorts. (F) to (G). Kaplan-Meier curves showing that patients with higher infiltration of POSTN ^high^ fibroblasts (F), and SPP1 ^high^ macrophages (G) are associated with worse outcomes. (H). Spatial plots of POSTN ^high^ fibroblasts (left) and SPP1 ^high^ macrophages (middle) identified using BayesSpace, alongside scatter plots showing the spatial correlation between POSTN ^high^ fibroblasts and SPP1 ^high^ macrophages in spatial transcriptomics (ST). (I). GSEA of 10 hallmark pathways between POSTN/SPP1 ^high^ and POSTN/SPP1 ^low^ groups in TCGA-PAAD cohort, with genes ranked by fold change in expression between the two groups. NES, normalized enrichment score.

Then, based on the expression levels of POSTN and SPP1, the TCGA-PAAD cohort was divided into two groups: POSTN ^high^ SPP1 ^high^ and POSTN ^low^ SPP1 ^low^. Hallmark gene set enrichment analysis (GSEA) was performed for these groups (Fig. 3I). The POSTN ^high^ SPP1 ^high^ group was enriched in tumor microenvironment-related pathways such as angiogenesis, hypoxia, and coagulation, as well as tumor-related epithelial-mesenchymal transition (EMT), the P53 pathway, and Notch signaling. Additionally, immune and inflammatory response-related pathways, including the IL2 STAT5 signaling, IL6 JAK STAT3 signaling, and complement pathways, were also enriched in this group. Within the fibroblast and macrophage/monocyte subpopulations, the POSTN ^high^ Fibro cluster was associated with hallmark pathways such as EMT and invasion (Fig. S6A). Similarly, the SPP1 ^high^ Macro cluster was linked to hallmark pathways, including EMT, invasion, metastasis, the cell cycle, and DNA repair (Fig. S6B). Cell states are typically regulated by transcription factor elements. In the POSTN ^high^ Fibro cluster, activated transcription factor regulons included GATA4 and BACH1 (Fig. S6C). The transcription factor BACH1 is activated under reduced oxidative stress conditions and stimulates lung cancer metastasis [37]. In the SPP1 ^high^ Macro cluster, the activated transcription factor regulons included FOXL1, FOXJ1, CEBPE, ISL1, and IRF6 (Fig. S6D). The CEBPE protein is a critical transcription factor required for the function and development of macrophages [38]. Overall, these results reveal the interaction between POSTN ^high^ Fibro and SPP1 ^high^ Macro within the PDAC TME and their role in promoting PDAC progression.

### CCL4 ^high^ CD8+ effector T cells and IGHG1 ^high^ IgG plasma cells are associated with favorable prognosis in PDAC patients

In the PDAC microenvironment, major cell types, such as T/NK and B/Plasma cells exhibited strong correlations across 12 independent bulk RNA-seq cohorts (Fig. 2B, and 2E). Tumor-infiltrating B cells enhance T-cell mediated antitumor immunity [39]. Antigen processing and presentation are fundamental to immunity, including antibody-dependent cellular cytotoxicity processes [39, 40]. Next, we subdivided and identified the subgroups of T/NK and B/Plasma cells. The subset of CD3-positive (CD3D or CD3E high) cells was used for batch correction and re-dimension reduction clustering. Fourteen clusters were delineated from T/NK cells (Fig. S7A). T cells and NK cells were classified on the basis of characteristic genes such as T cells (CD3, CD4, and CD8) and NK cells (FCGR3A, NCAM1). The expression of regulatory (FOXP3), naive (LEF1 and SELL), central memory (IL7R, CCR7, TCF7, and LTB), effector (named eff; IFNG), effector memory (named EM; GZMK, PRF1, GZMB, and GNLY), costimulatory (CD27, CD28, TNFRSF4, TNFRSF9, and TNFRSF18), and exhausted (PDCD1, HAVCR2, LAG3, LAYN, TIGIT, and ENTPD1) markers was subsequently used to define the 14 cell clusters (Fig. S7B). Each cluster’s representative gene was used to define the final cluster names, including LTB ^high^ CD4+ T _Naive_, GZMK ^high^ CD8+ T_EM_, HSPA1A ^high^ T _stressed CM_, GNLY ^high^ NK, MT1E ^high^ CD8+ T, CCL4 ^high^ CD8+ T_EFF_, BATF ^high^ T _Reg_, FOS ^high^ CD8+ T_CM_, CXCL13 ^high^ CD4+ T _EM_, XCL1 ^high^ NK, CCL20 ^high^ CD4+ T_CM_, CH25H^high^ CD4+ T _Naive_, and MKI67 ^high^ CD8+ T _EM_ (Fig. S7E). Interestingly, T/NK cell infiltration, such as LTB ^high^ CD4+ T _Naive_, BATF ^high^ T _reg_, GZMK ^high^ CD8+ T _EM_, and MKI67 ^high^ CD8+ T _EM_, was more pronounced in tumor tissues than in NP and AT sites (Fig. 3A and 3B). In the LM sites, the T/NK cell subsets displayed infiltration of central memory T cells, including HSPA1A ^high^ CD8+ T _stressed CM_ and MT1E ^high^ CD8+ T_CM_. In contrast, ATs presented a greater proportion of LTB ^high^ CD4+ T _Naive_ cells (Fig. 4B). And some GZMK ^high^ CD8+ T cells, CCL4 ^high^ CD8+ T cells, GNLY ^high^ NK cells, and a small proportion of BATF ^high^ T _Reg_ cells enriched in Ats (Fig. 4A). Notably, the proportion of BATF ^high^ T _Reg_ infiltration was higher in PDAC primary tissues than in LM, AT, and NP tissues (Fig. 4B). Moreover, BATF ^high^ T _Reg_ cells demonstrated costimulatory functions and expressed exhaustion markers such as TIGIT, LAYN, and ENTPD1 (Fig. S7B).

**Figure 4.**
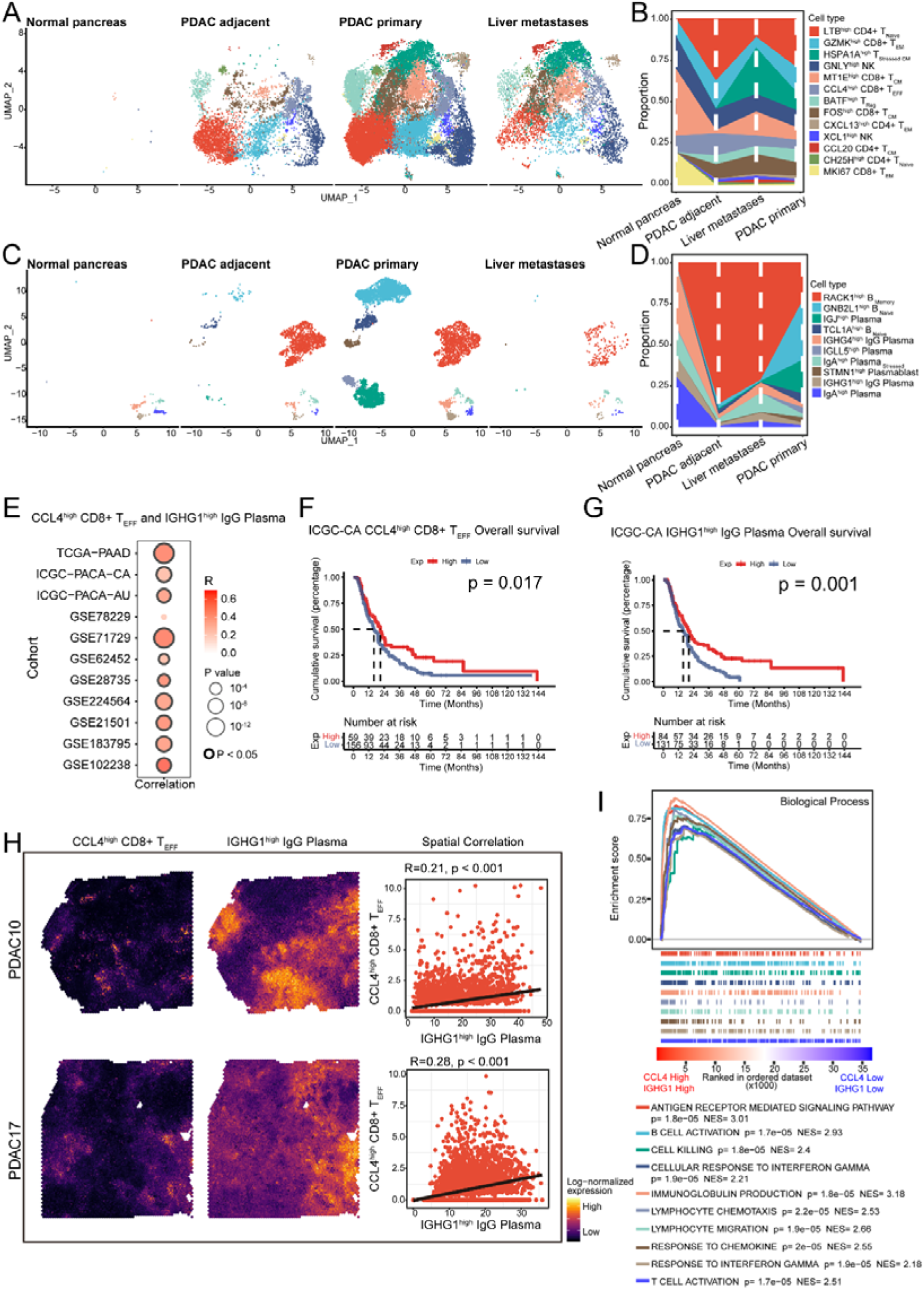
Decoding the Immune Signature of CCL4 ^high^ CD8+ effector T and IGHG1 ^high^ IgG Plasma cells in PDAC TME. (A). UMAP plots showing the composition of T/NK colored by cell subtype. (B). The percentage of each T or NK cell subtypes in different tissue sites in the scRNA-seq data. (C). UMAP plots displaying the composition of B/Plasma cells colored by subtype. (D). The percentage of each B or Plasma cell subtypes across different tissue sites in scRNA-seq data. (E). A dot plot illustrates the Spearman correlation coefficients (R) and their corresponding p-values between CCL4 ^high^ CD8+ T cells and IGHG1 ^high^ plasma cells across 11 independent cohorts. (F) to (G). The Kaplan-Meier curves show that patients with higher infiltration of CCL4 ^high^ CD8+ T_EFF_ (F), and IGHG1 ^high^ IgG Plasma (G) are associated with better outcomes. (H). Spatial plots of CCL4 ^high^ CD8+ T_EFF_ (left) and IGHG1 ^high^ IgG Plasma (mid) using BayesSpace, alongside scatter plots of the spatial correlation between CCL4 ^high^ CD8+ T_EFF_ and IGHG1 ^high^ IgG Plasma in spatial transcriptomics (ST). (I). GSEA of 10 GO BP terms comparing the CCL4 & IGHG1 high and CCL4 & IGHG1 low groups in the TCGA-PAAD cohort. Genes are ranked by fold change in expression between these two groups. NES, normalized enrichment score. BP, biological process.

Similarly, we subsetted the B/Plasma cells and performed re-dimensional clustering into 10 sub-clusters. Based on the expression of MS4A1, CD19, CD38, JCHAIN, MZB1, and SDC1, the cells were classified as B cells, plasma cells, or plasmablasts. In accordance with the findings of Hao *et al*. on B cells and plasma cells [41], we categorized the cell states using markers as naive (SELL, IL4R, FCER2), memory (CD24, CD27), stressed (BAG3, HSPA6, HSPB1), proliferative (MKI67, STMN1), IgG (IGHG1, IGHG2, IGHG3, IGHG4), IgA (IGHA1, IGHA2), IgM (IGHM), and IgD (IGHD). The 10 subclusters were defined and named as RACK1 ^high^ B memory, GNB2L1 ^high^ B _naive_, TCL1A ^high^ B_naïve_, IGJ ^high^ Plasma, IGHG4 ^high^ IgG Plasma, IGLL5 ^high^ Plasma, IgA ^high^ Plasma _stressed_, IGHG1^high^ IgG Plasma, IgA ^high^ Plasma, and STMN1^high^ Plasmablast (Fig. 4C and 4D). Notably, the infiltration of B/Plasma subclusters among different tissues was inconsistent, with PDAC primary tissues infiltrated by all subtypes (Fig. 4C and 4D). And plasma cells expressing IgA or IgG antibodies were present across all four tissue origins (Fig. 4C). The enrichment of IgG and IgA plasma cells may contribute to an immune ecology that aids in infection defense[42]. Furthermore, PDAC primary sites were infiltrated by B naive cells, including GNB2L1 ^high^ B _naive_ and TCL1A ^high^ Plasma. Additionally, compared with the NP, AT, and LM sites, PDAC infiltrated two types of plasma cells: IGJ ^high^ plasma cells and IGLL5 ^high^ plasma cells. These cells were characterized by high expression of memory gene CD27 and low expression of IgA, IgG, IgM, or IgD molecules (Fig. S7D). This part of the study comprehensively compared the characteristics of immune cell infiltration among PDAC, LM, NP, and AT.

We subsequently performed a correlation analysis of the GSVA enrichment scores for representative genes of different cell types. The results revealed strong correlations between LTB ^high^ CD4+ T _Naive_ and RACK1 ^high^ B _memory_, as well as GNB2L1 ^high^ B _naive_ cells across 12 bulk RNA-seq cohorts (Fig. 4E). However, high scores for LTB ^high^ CD4+ T _Naive_, RACK1 ^high^ B _memory_, and GNB2L1 ^high^ B _naive_ individuals were associated with poor or insignificant patient prognosis (Fig. S8). Additionally, there was a strong correlation between FOS ^high^ CD8+ T _CM_ and TCL1A ^high^ B _naive_, as well as between MKI67 ^high^ CD8+ T _EM_ and TCL1A ^high^ B _naive_, and STMN1 ^high^ Plasmablast (Fig. S9A). High scores for these cells were associated with poor or insignificant patient prognosis (Fig. S8 and S9A). Additionally, MT1E ^high^ CD8+ T _CM_ and CXCL13 ^high^ CD4+ T _EM_ were positively correlated with IgA ^high^ Plasma. High gsva scores of IgA ^high^ Plasma were associated with favorable prognosis in the ICGC-CA cohort (p=0.01), but not significant in the ICGC-AU cohort (p=0.153). High MT1E ^high^ CD8+ T _CM_ scores were unfavorable factors in the ICGC-AU cohort (p=0.019). High scores of CXCL13 ^high^ CD4+ T _EM_ were favorable in the ICGC-CA cohort (p=0.003), but not significant in the ICGC-AU cohort (p=0.117). CD4+ T cells expressing CXCL13 can recognize tumor neoantigens and are categorized as memory T cells, which are also associated with CD8+ T cell status and patient survival, as reported in another melanoma study [43].

Although most T/NK subgroups and B/Plasma subgroups related to patient prognosis presented unfavorable relationships, CCL4 ^high^ CD8+ T _EFF_ and IGHG1 ^high^ IgG plasma levels were strongly correlated, with correlation coefficients ranging from 0.32 to 0.64 across 10 datasets (p<0.05), whereas only 0.25 in the GSE78229 dataset (p=0.08; Fig. 4E and S9A). High infiltration scores of CCL4 ^high^ CD8+ T _EFF_ cells (p=0.017) and IGHG1 ^high^ IgG plasma cells (p=0.001) were favorable prognostic factors in the ICGC-CA cohort (Fig. 4F and 4G). Similarly, in the ICGC-AU cohort, high cell scores of CCL4 ^high^ CD8+ T _EFF_ cells (p=0.06) and IGHG1 ^high^ IgG plasma cells (p=0.001) were favorable for patient prognosis (Fig. S9B and S9C). Furthermore, the spatial colocalization of these two cell types CCL4 ^high^ CD8+ T _EFF_ cells and IGHG1 ^high^ IgG plasma cells was observed, with the spatial correlation of CCL4 ^high^ CD8+ T _EFF_ cells and IGHG1 ^high^ IgG plasma cells ranging from 0.1 to 0.28 in 11 ST-seq datasets (Fig. 4H and S9D). The TCGA-PAAD samples were divided into two groups based on the simultaneous high or low expression of CCL4 and IGHG1 for Gene Ontology Biological Process (GO BP) GSEA. The CCL4 ^high^ IGHG1 ^high^ group was significantly enriched in processes related to T cell activation, B cell activation, antigen receptor-mediated signaling, the cellular response to interferon-gamma, and the cell killing function (Fig. 4I). These results reveal the immunosuppressive nature of the PDAC tumor microenvironment and suggest that the positive correlation between CCL4 ^high^ CD8+ T _EFF_ cells and IGHG1 ^high^ IgG plasma cells may improve patient prognosis.

### Expression of POSTN, SPP1, CCL4, and IGHG1 and their potential as therapeutic targets

POSTN was expressed mainly in fibroblasts within tumor tissues, including those of primary PDAC tumors and liver metastases (Fig. 5A). SPP1 was also expressed in tumor tissues, specifically in Macro/Mono cells from PDAC primary and LM patients, although notably, SPP1 was detected in epithelial and endothelial cells (Fig. 5A). Moreover, the mRNA expression levels of POSTN and SPP1 were lower in non-tumor regions than in tumor areas (Fig. 5A). High expression of POSTN (p=0.009) and SPP1 (p<0.001) was associated with poor prognosis, with patients with high levels of both markers (p<0.001) experiencing even worse outcomes (Fig. 5B). Furthermore, analysis of a combined bulk RNA dataset revealed a strong correlation between POSTN and SPP1 expression (R=0.87, p<0.001; Fig. 5C). Multiplex immunohistochemistry (mIHC) of human PDAC tissues confirmed that POSTN and SPP1 colocalized with a fibroblast marker (α-SMA) and a macrophage marker (CD163), with both cell types clustering near the tumor marker (CK19; indicated by white/yellow arrows; Fig. 6A). Notably, the co-localization of the tumor marker CK19 and the fibroblast marker α-SMA may be related to the presence of POSTN ^high^ fibroblasts, which are CAFs. CAFs express both CK19 and α-SMA in the TME [44]. POSTN, a key extracellular matrix protein secreted by fibroblasts, particularly CAFs, is likely released into the tumor stroma, where it interacts with tumor cells to promote tumor growth and invasion[45]. Additionally, the co-expression of POSTN and SPP1, along with their high correlation in tumor tissues, suggests a synergistic effect of these factors in the TME (Fig. 5C and 6A). This interaction may enhance PDAC invasiveness by promoting the activity of POSTN ^high^ CAFs and SPP1 ^high^ macrophages, thereby accelerating tumor progression.

**Figure 5.**
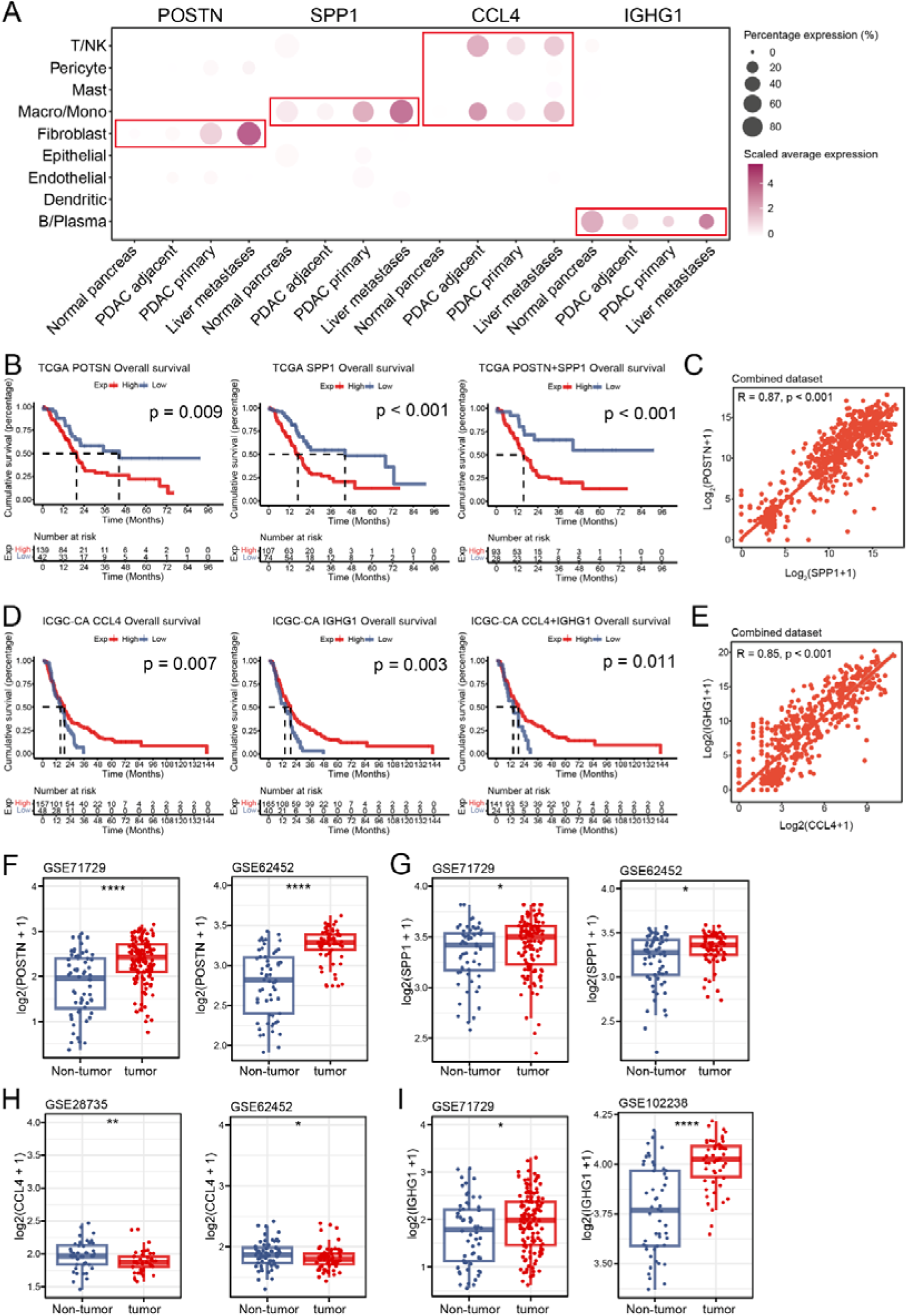
Gene expression of POSTN, SPP1, CCL4, and IGHG1 at the single-cell level and bulk RNA level across multiple independent cohorts. (A). The bubble plot shows the expression levels of POSTN, SPP1, CCL4, and IGHG1 across different cell types (left) and across various tissue sites (bottom), with average expression scaled by SCTransform. (B-C). The box plot shows the comparison of POSTN (C) and SPP1 (D) expression between non-tumor and tumor samples in the bulk RNA-seq datasets GSE71729 and GSE62452. Expression values are scaled by log2(expression+1). Statistical significance was determined using the Wilcoxon test. (D-E). The box plot shows the comparison of IGHG1 expression between non-tumor and tumor samples in the bulk RNA-seq datasets GSE71729 and GSE102238. Expression values are scaled by log2(expression + 1). Statistical significance was determined using the Wilcoxon test. (F) The Kaplan-Meier curves show that patients with higher expression of POSTN (left), SPP1 (middle), and both POSTN & SPP1 (right) are associated with worse prognosis in TCGA dataset. (G) Dot plot shows the positive correlation between POSTN and SPP1 expression in the combined dataset. The expression values are scaled as log2(expression + 1). (H) The Kaplan-Meier curves show that patients with higher expression of CCL4 (left), IGHG1 (middle), and both CCL4 & IGHG1 (right) are associated with better prognosis in ICGC-CA cohort. (I) The dot plot shows the positive correlation between CCL4 and IGHG1 expression in the combined dataset. The expression values are scaled as log2(expression + 1).

**Figure 6.**
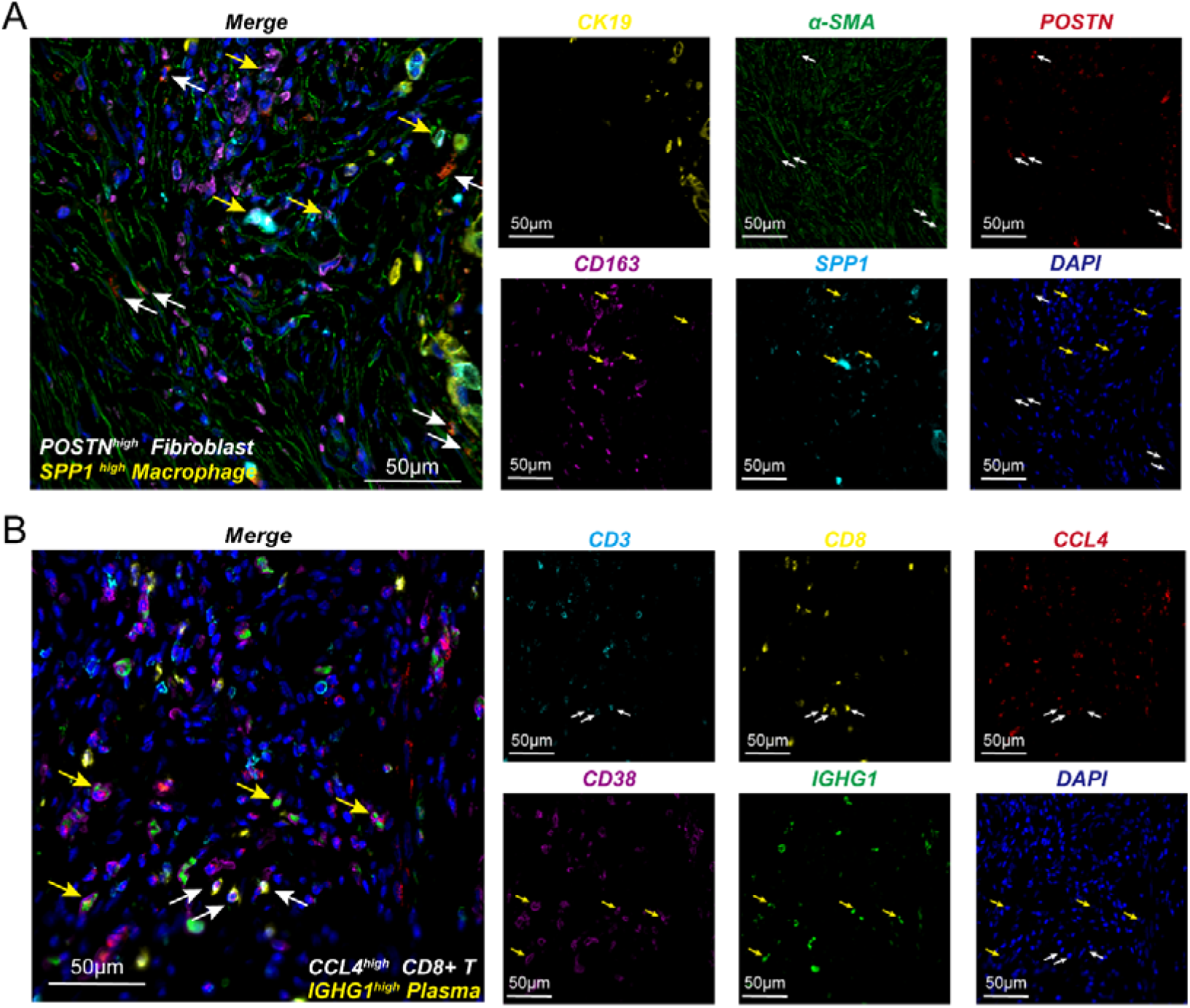
Validation of POSTN ^high^ fibroblasts, SPP1 ^high^ macrophages, CCL4 ^high^ CD8+ T cells, and IGHG1 ^high^ plasma cells in human PDAC tissues via mIHC. (A). Representative immunofluorescence (IF) staining of human PDAC tissue. The merge, DAPI (blue), CK19 (yellow), α-SMA (green), POSTN (red), CD163 (purple), and SPP1 (light blue) are shown in individual and merged channels. Bar, 50 μm. The experiment was performed on one sample from PDAC patient. The white arrows indicate the POSTN ^high^ fibroblasts, and the yellow arrows indicate the SPP1 ^high^ macrophages. (B). Representative IF staining of human PDAC tissue. The merge, DAPI (blue), CD3 (bright blue), CD8 (yellow), CCL4 (red), CD38 (purple), and IGHG1 (green), are shown in individual and merged channels. Bar, 50 μm. The experiment was performed on another sample from PDAC patient. The white arrows indicate CCL4^high^ CD8+ T cells, while yellow arrows indicate IGHG1^high^ plasma cells.

In our study, CCL4 was highly expressed in T cells at the single-cell level, primarily in tumors or PDAC adjacent regions (Fig. 5A). CCL4 plays a key role in recruiting cytotoxic CD8+ T cells into the tumor microenvironment, potentially enhancing immune responses against PDAC[46, 47]. The mIHC results revealed that CCL4 co-localized with CD8-positive T cells (CD3 and CD8) in the tumor microenvironment (indicated by white arrows; Fig. 6B). This aligns with our single-cell analysis, where CCL4 ^high^ CD8+ T cells were present in PDAC tissues, albeit in small proportion (Fig 4B). More importantly, the presence of CCL4 ^high^ CD8+ T cells was significantly associated with better prognosis in PDAC patients, contrasting with other immune cell populations that showed a correlation with worse prognosis (Fig S8). This suggests that CCL4 ^high^ CD8+ T cells play a protective role in PDAC by promoting anti-tumor immune responses. Interestingly, we found that CCL4 expression was greater in non-tumor regions than in tumor areas (Fig. 5H), which may indicate that cytotoxic CD8+ T cells accumulate in PDAC adjacent areas but have difficulty effectively infiltrating primary tumor regions in the PDAC microenvironment. This observation could be due to the immunosuppressive nature of the PDAC stroma, which hinders effective immune cell infiltration. On the other hand, IGHG1, predominantly expressed by plasma cells, was highly expressed in tumor regions and partially infiltrated liver metastasis sites (Fig. 5A). Although this expression pattern was somewhat inconsistent with the single-cell results (Fig. 5A and 5I), the high expression of CCL4 and IGHG1 was still associated with favorable patient prognosis (Fig. 5D). In the combined bulk RNA dataset, CCL4 and IGHG1 expression showed a strong correlation (R=0.85, p<0.001; Fig. 5E), supporting their potential interplay within the TME. Furthermore, the mIHC staining also revealed co-localization of a plasma cell marker (CD38) with IGHG1 in the PDAC tumor microenvironment (indicated by yellow arrows; Fig 6B). CCL4 co-localized with CD8-positive T cells (CD3 and CD8), and both cell types were found to cluster in close proximity to each other (Fig. 6B). Notably, we observed that CCL4 does not fully colocalize with CD8+ T cells in the PDAC TME. This discrepancy may arise because, although CCL4 is expressed on CD8+ T cells, its primary role in the tumor microenvironment is likely to be mediated through secretion, which recruits additional CD8+ T cells to the tumor site[48]. However, these findings suggest that CCL4 ^high^ CD8+ T cells and IGHG1 ^high^ IgG plasma cells are spatially close in the TME, which may enhance their cooperative roles in promoting anti-tumor immunity in PDAC. Taken together, these results validate our hypothesis that the presence of CCL4 ^high^ CD8+ T cells and IGHG1 ^high^ IgG plasma cells within the TME is associated with improved prognosis in PDAC patients.

### Spatial mapping of POSTN ^high^ fibroblasts, CCL4 ^high^ CD8+ T Cells, and IGHG1 ^high^ IgG plasma cells within PDAC using Xenium high-resolution spatial transcriptomics

Pancreatic intraepithelial neoplasia (PanIN) is the most common and well-studied precursor lesion. To further investigate and validate whether POSTN ^high^ fibroblasts play a carcinogenic role in the early PDAC TME, we analyzed a sample from a PDAC surgical specimen with associated high-grade PanIN[33]. After standard data filtering and cell dimensionality reduction clustering, nine clusters were identified (Fig. S10A). Using cell type markers and referencing PanIN markers, the spatial regions were categorized into PanIN, Normal, Stroma, Immune, and Mixed regions (Fig. S10B). Based on POSTN expression, POSTN ^high^ Fibro & Macro and Fibro & Macro regions (Low expressed level of POSTN) were further differentiated within the Stroma and Immune regions (Fig. 7A). POSTN expression was primarily concentrated in the POSTN ^high^ Fibro & Macro cluster, with minimal expression observed in the PanIN region (Fig. S10C). Focusing on the area surrounding the PanIN region, we observed a significant accumulation of POSTN ^high^ fibroblasts and macrophages around the boundaries of high-grade PanIN lesions (Fig. 7B). Two spatial areas were selected, with the clearly visible PanIN regions outlined in bright yellow (Fig. 7C and S11A). Using the single-cell resolution of the Xenium platform, we identified fibroblasts in the PanIN microenvironment that simultaneously expressed POSTN and COL1A1 within the cell segmentation defined by DAPI staining (Fig. 7C and S11A). These POSTN ^high^ fibroblast cells were primarily located at the boundaries and periphery of PanIN region, where TFF1 was expressed (Fig. 7C). Due to the absence of SPP1 detection in this sample, we could only observe the expression of macrophages (CD163), which were also notably clustered around the edges of the PanIN region. Similarly, in Spatial Area 2, POSTN ^high^ fibroblasts were observed to accumulate at the boundaries and periphery of the PanIN lesions (Fig. S11A). In both spatial areas, the proximity or overlap between POSTN ^high^ fibroblasts and macrophages was evident, suggesting their combined role in influencing the progression of PanIN. These findings indicate that POSTN ^high^ fibroblasts influence the progression of high-grade PanIN, indirectly supporting the carcinogenic role of this cell subtype in the early PDAC TME.

**Figure 7.**
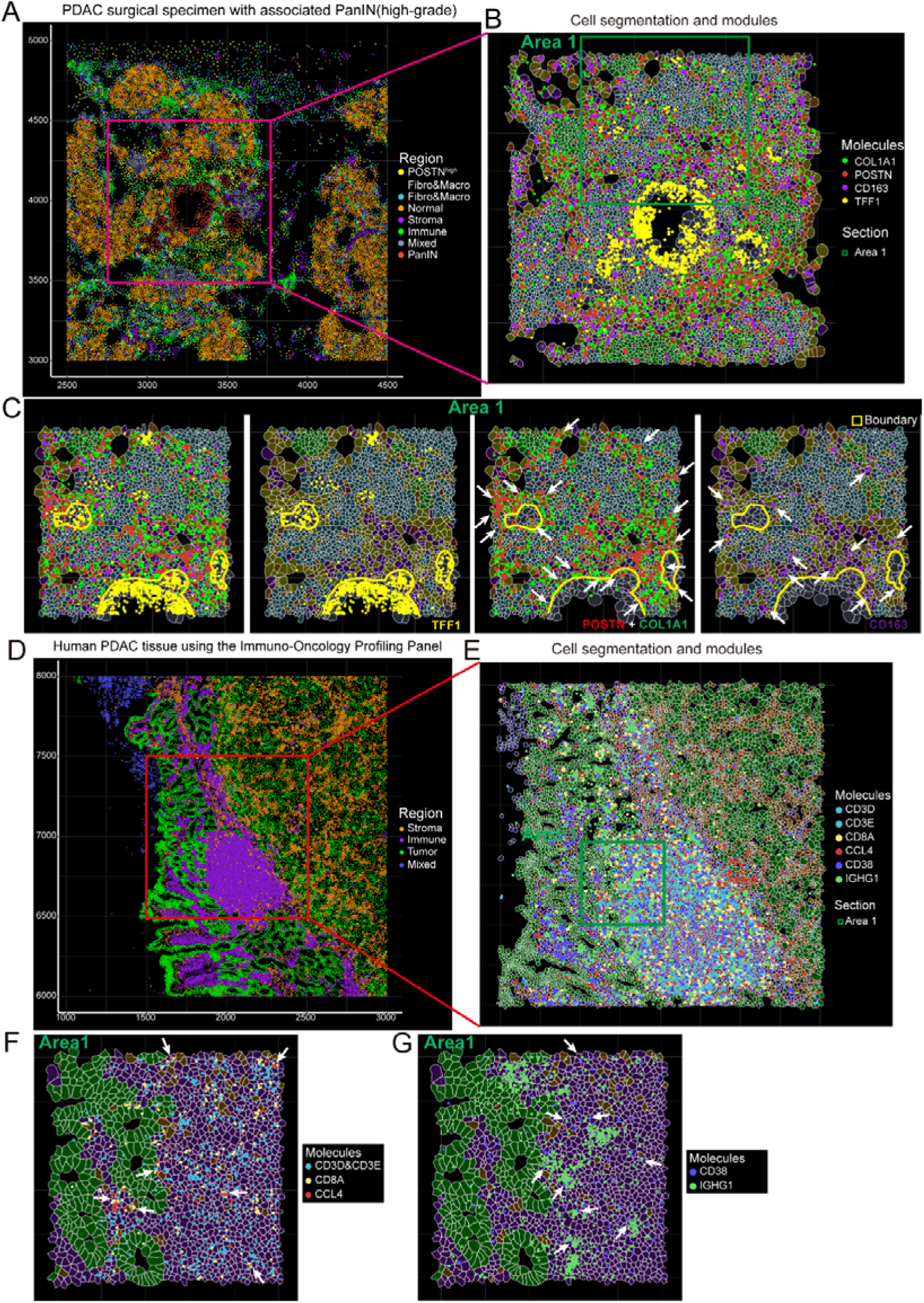
High-resolution spatial omics characterization of POSTN ^high^ fibroblasts in PDAC with high-grade PanIN lesions, and characterization of CCL4 ^high^ CD8+ T cells and IGHG1 ^high^ plasma cells in a Xenium platform with an immune-oncology panel. (A) Spatial region distribution map of PDAC surgical specimen with associated PanIN (high-grade) via Seurat V5. (B) Spatial distribution of cell segmentation and associated modules in high-grade PanIN tissue. (C) The spatial distribution maps of cell segmentation and modules in Area1 show Merge, TFF1 (yellow), POSTN+COL1A1(red+green), and CD163(purple), respectively. (D) An annotated spatial region distribution map of human PDAC tissue, generated using an Immuno-Oncology Profiling Panel provided by the 10x Genomics dataset. (E) Spatial distribution of cell segmentation and associated modules in PDAC TME. (F) The spatial distribution maps of cell segmentation and modules in Area1 show CD3D&CD3E (blue), CD8A (yellow), and CCL4 (red), respectively. (G) A spatial distribution maps of cell segmentation and modules in Area1 show CD38 (purple), and IGHG1 (green), respectively.

The inclusion of an immune-target panel in the PDAC RNA in situ hybridization sample is promising. We utilized this sample to further investigate whether CCL4-expressing CD8+ T cells and IGHG1-secreting plasma cells are present in the PDAC TME. Using the same standard processing applied to the high-grade PanIN sample, we identified 8 spatial clusters (Fig. S10D). Due to the different gene types included in the panels, we characterized the Immune region using markers for T cells (CD3D, CD3E), macrophages (CD68), plasma cells (CD38), and NK cells (NKG7); the Stroma region was delineated by ACTA2 and DCN; and the Tumor region was identified by EPCAM (Fig. S10E). Subsequently, Stroma, Immune, Tumor, and Mixed regions were identified (Fig. 8A). We selected an area enriched in immune cells for further analysis of CCL4 ^high^ CD8+ T cells and IGHG1 ^high^ plasma cells (outlined in red; Fig. 8A). After mapping the cell segmentation and associated modules (including CD3D, CD3E, CD8A, CCL4, CD38, and IGHG1), three spatial areas were selected for further characterization (Fig. 7D and S11B). We defined CCL4 ^high^ CD8+ T cells in the PDAC TME as cells that simultaneously express CD3D or CD3E, CD8A, and CCL4. The identification method for IGHG1 ^high^ plasma cells was similar, requiring the simultaneous expression of IGHG1 and CD38 in a single cell. The results were exciting: in all three spatial areas, CCL4 ^high^ CD8+ T cells were present (indicated by white arrows; Fig. 7F and S11C). Furthermore, plasma cells expressing IGHG1 were also observed in these areas (indicated by white arrows; Fig. 7G and S11D). By comparing the spatial distribution of CCL4 ^high^ CD8+ T cells and IGHG1 ^high^ IgG plasma cells, we found that some of these two cell types were found in close proximity. These results suggest the presence of CCL4 ^high^ CD8+ T cells and IGHG1 ^high^ plasma cells in the PDAC TME.

In summary, our findings highlight that POSTN and SPP1 may serve as potential therapeutic targets within the PDAC microenvironment. Notably, promoting the infiltration of CCL4 ^high^ CD8+ T cells and IGHG1 ^high^ IgG plasma cells, along with modulating the interaction between POSTN ^high^ fibroblasts and SPP1 ^high^ macrophages, could provide novel therapeutic strategies for improving patient outcomes in PDAC by influencing tumor progression through the complex TME interactions.

## Discussion

The advent of scRNA-seq and ST-seq has profoundly expanded our understanding of tumor biology, enabling an unprecedented level of detail in the characterization of the TME [17, 49, 50]. These technologies have allowed researchers to dissect the cellular and molecular composition of the TME with unparalleled precision, revealing the intricate interactions that govern tumor progression and resistance to therapy [51, 52]. However, despite these advancements, the cellular architecture and intercellular interactions within the PDAC microenvironment remain incompletely understood. In this study, we conducted a comprehensive integrative analysis to map the cellular landscape of the human PDAC TME, utilizing a robust dataset comprising 88 PDAC-related single-cell studies.

Our analysis provides a detailed overview of the cellular constituents of the PDAC TME, revealing significant variations in cell type distribution across tumor, adjacent, and metastatic tissues. Notably, we observed a marked enrichment of fibroblasts within primary PDAC tissues, with a significantly higher proportion than in LM tissues and a trend toward higher levels than in NP and AT tissues. These findings corroborate previous studies that have implicated CAFs in promoting PDAC progression through extracellular matrix remodeling, modulating immune infiltration, and secreting pro-tumorigenic factors [53, 54]. Additionally, macrophages were found to be more abundant in primary PDAC tissues than in NP tissues, further underscoring their role in shaping a pro-tumorigenic microenvironment. Conversely, the infiltration of immune cells, particularly T cells and NK cells, was predominantly localized to adjacent tissues, with significantly lower levels observed within the tumor core. This spatial distribution is likely attributable to the recruitment of immunosuppressive myeloid cells and the establishment of a CAF-rich stroma, which collectively act as physical and biochemical barriers to immune cell infiltration, particularly that of CD8+ T cells [29, 55, 56]. Our findings highlight the spatial heterogeneity of the PDAC TME, with fibroblasts and macrophages concentrated within the tumor core and immune cells predominantly localized to peritumoral regions.

Intercellular interactions within the TME are critical determinants of tumor initiation, progression, and immune evasion. Through integrative scRNA-seq and bulk RNA-seq analyses, we identified fibroblasts and macrophages/monocytes as the predominant interacting cell types within the PDAC TME, with additional interactions observed between B/plasma cells and T/NK cells. Our spatial transcriptomics and mIHC analyses further revealed significant colocalization of POSTN ^high^ fibroblasts and SPP1 ^high^ macrophages within tumor tissues. POSTN, an extracellular matrix protein secreted by stromal cells, has been implicated in enhancing the activity of M2 macrophages and CAFs through the integrin-mediated NF-κB and TGF-β2 signaling pathways, thereby promoting metastatic dissemination in various malignancies, including ovarian cancer [57, 58]. Similarly, SPP1, a key mediator of TAMs function, was found to be highly expressed in macrophages within both primary PDAC and LM tissues. SPP1 has been recognized as a pivotal factor in promoting metastatic colonization, particularly in colorectal cancer [59], where SPP1 ^high^ macrophages, in conjunction with FAP ^high^ fibroblasts, orchestrate the fibrotic architecture of the TME [60]. The spatial colocalization of POSTN ^high^ fibroblasts and SPP1 ^high^ macrophages observed in our study suggests that their interaction may play a critical role in establishing a hypoxic, TNFα-rich microenvironment conducive to tumor progression through NF-κB signaling.

PDAC is widely recognized as an “immunologically cold” tumor characterized by a highly immunosuppressive TME that poses significant challenges to the efficacy of immune-based therapies. Our study identified several immune cell populations within the PDAC TME, including CD4+ T cells expressing high levels of BATF, LTB/CH25H, and CCL20, corresponding to regulatory T cells (Tregs), naïve T cells, and central memory T cells (TCMs), respectively. Tregs have been extensively documented as key players in promoting immunosuppression and sustaining chronic inflammation within the PDAC TME, yet their presence does not appear to significantly impact patient prognosis [61]. In contrast, our data revealed that CCL4 ^high^ CD8+ T cells are associated with improved patient outcomes, despite their relatively low infiltration levels. The chemokine CCL4 is known to facilitate the recruitment and activation of CD8+ T cells, suggesting a potential role in enhancing the cytotoxic capacity of these cells within the TME [47, 62]. In the pancreatic cancer TME, chemokines, including CCL4, are closely associated with the infiltration and activation of CD8+ T cells[62]. Another study has shown that CCL4 is highly expressed in infiltrating CD8+ effector T cells within the TME of non-small cell lung cancer (NSCLC), which is consistent with our findings [47]. Furthermore, the presence of IGHG1-secreting plasma cells was positively correlated with favorable prognosis, particularly in patients with concomitant high expression of IGHG1 and CCL4. These patients exhibited a TME enriched with pathways related to immune activation, including T-cell activation, lymphocyte migration, cytotoxicity, and the chemokine response. Spatial analyses further confirmed the close proximity of CCL4 ^high^ effector CD8+ T cells and IGHG1 ^high^ IgG plasma cells, implying a potential synergistic interaction that may potentiate CD8+ T-cell mediated cytotoxicity through IGHG1 secretion by plasma cells. That, our findings highlight the potential of CCL4 ^high^ CD8+ T cells and IGHG1-secreting plasma cells in enhancing immune responses within the PDAC TME, suggesting a synergistic interaction that could be leveraged to improve immunotherapy outcomes.

While our study provides novel insights into the cellular composition and intercellular interactions within the PDAC TME, several limitations warrant consideration. We used a PDAC surgical sample with high-grade PanIN lesions in combination with a Xenium sample to validate the spatial localization of POSTN-high fibroblasts in the PanIN-to-carcinoma transition. While this indirectly supports the role of these cells in the early PDAC TME, future validation with appropriate PDAC tumor samples would provide more compelling evidence. And the absence of functional validation experiments, particularly in vivo and in vitro assays, limits the direct translation of our findings into therapeutic strategies. Future studies should prioritize experimental validation to corroborate the interactions and functional roles of the identified cell populations within the PDAC TME.

Nevertheless, our integrative analysis offers a comprehensive overview of the PDAC TME, identifying critical cellular interactions that may drive tumor progression and immune evasion, particularly the interplay between POSTN ^high^ fibroblasts and SPP1 ^high^ macrophages, and the potential immunostimulatory role of CCL4 ^high^ CD8+ T cells in conjunction with IGHG1-secreting plasma cells. These findings provide a foundation for the development of targeted therapeutic approaches aimed at modulating the TME to enhance anti-tumor immunity and improve outcomes in patients with PDAC.

## Conclusion

This study provides a comprehensive analysis of the PDAC tumor microenvironment, revealing key interactions between POSTN ^high^ fibroblasts and SPP1 ^high^ macrophages that create a pro-tumorigenic niche, while identifying the beneficial role of CCL4 ^high^ CD8+ T _EFF_ cells and IGHG1 ^high^ IgG plasma cells in patient prognosis. These findings suggest that targeting these cellular interactions could offer new therapeutic strategies to enhance immune responses and improve outcomes in PDAC patients.

## List of abbreviations

PDAC: Pancreatic ductal adenocarcinoma
TME: tumor microenvironment
scRNA-seq: single-cell RNA sequencing
ST-seq: spatial transcriptomics
GSVA: gene set variation analysis
mIHC: multiplex immunohistochemistry
CAFs: cancer-associated fibroblasts
ECM: extracellular matrix
TAMs: tumor-associated macrophages
LM: liver metastases
AT: PDAC adjacent-tumor
NP: normal pancreas
PCA: principal component analysis
UMAP: uniform manifold approximation and projection
DEGs: differential expressed genes
MSigDB: Molecular Signatures Database
GSEA: gene set enrichment analysis
GO BP: gene ontology biological process

## Declarations

### Ethics approval and consent to participate

This study was approved by the Ethics Committee of the Second Affiliated Hospital of Wannan Medical College, and was conducted in accordance with the Declaration of Helsinki (WYEFYLS2024101).

### Consent for publication

Not applicable.

### Availability of data and materials

The data and accession numbers used in this study can be found in the article and supplementary tables. The codes used for all processing and analysis are available at https://github.com/JunekureWu/integratedPDAC.

### Author contributions

Conceptualization: Qizhou Lian, Yang Liu, and Jierong Chen

Formal analysis: Jun Wu, Tenghui Dai, Ziyue Li, and Meng Pan

Visualization: Jun Wu, Meng Pan, and Tenghui Dai

Methodology: Meng Pan, Ziyue Li, and Wei Zhang

Data Curation: Hao Chen, Guansheng Zheng, and Li Qiao

Supervision: Qizhou Lian, Yang Liu, and Jierong Chen

Funding acquisition: Meng Pan, Qizhou Lian, and Jierong Chen

Writing - Original Draft: Jun Wu, Tenghui Dai, Ziyue Li, and Meng Pan

Writing - Review & editing: Jun Wu, Qizhou Lian, Yang Liu, and Jierong Chen

## Acknowledgements

The author J.W. sincerely acknowledges Ms. Y.C. for her invaluable support in this work.

