## Supplemental figures S1-S11 for "Integrating multi-modal transcriptomics identifies cellular subtypes with distinct roles in PDAC progression"

Jun Wu et al.

Here, as authors, we present the supplemental figures included in this study.

Supplementary Figures S1-S11 and their captions can be found in this document.

Figure S1

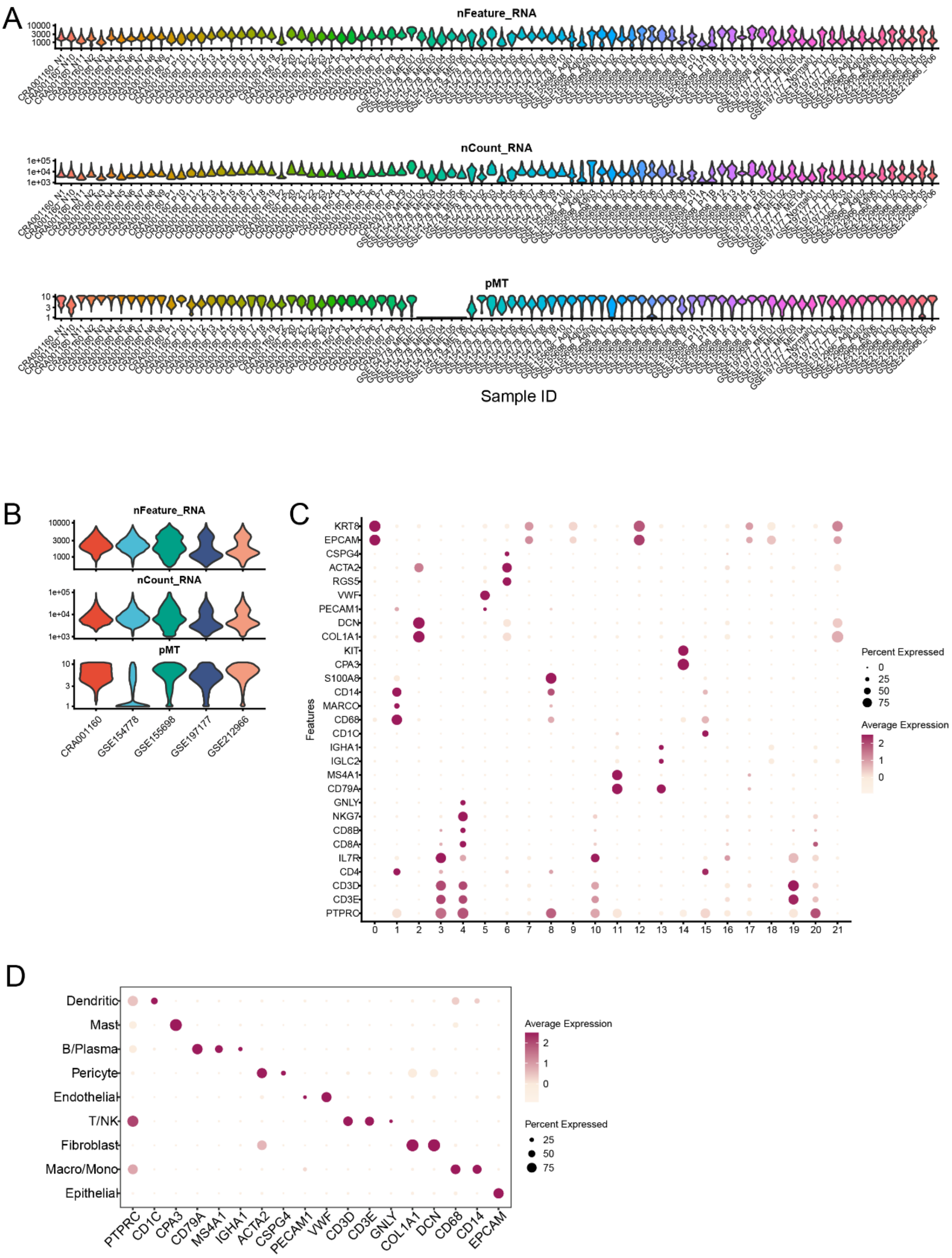


**Figure S1**. scRNA-seq cohort data quality control and cell type identification

A. Violin plots of Feature (nFeature_RNA), Count (nCount_RNA), and mitochondrial proportion (pMT) after quality control were generated for five cohorts: CRA001160 (n=35), GSE154778 (n=16), GSE155698 (n=20), GSE197177 (n=7), and GSE212966 (n=9).

B. Violin plots of the total Feature (nFeature_RNA), total Count (nCount_RNA), and total mitochondrial proportion (pMT) after quality control for the five cohorts were generated.

C. Bubble plot of the expression of cell markers for the 22 clusters generated by Seurat. The x-axis represents the clusters, and the y-axis represents the cell marker features.

D. Bubble plot of the expression of representative markers for each of the 9 major cell types. The x-axis represents the cell marker features, and the y-axis represents the cell types.

Figure S2 Spatial Atlas of PDAC

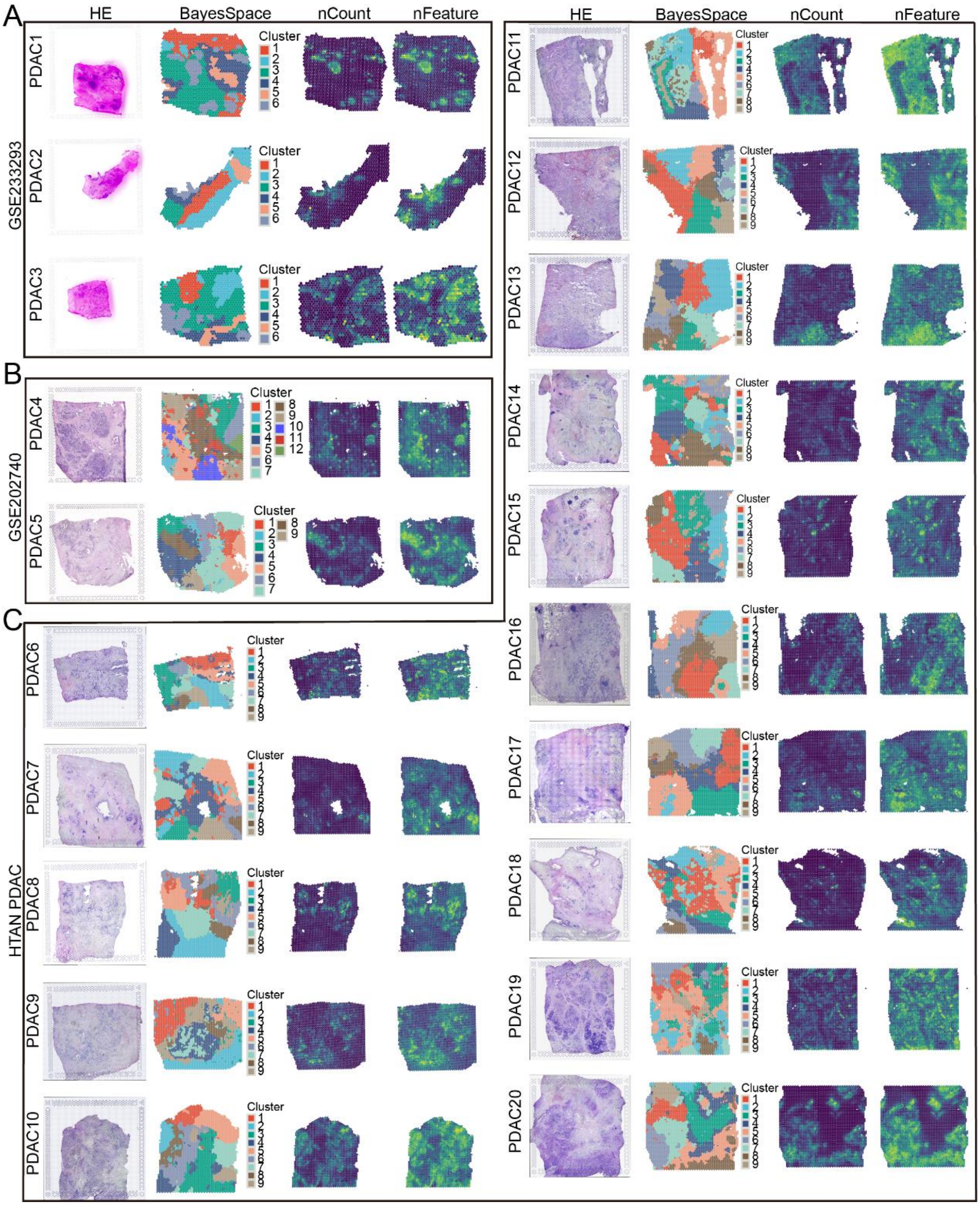


**Figure S2**. Quality control and dimensionality reduction clustering based on spots in spatial transcriptomics sequencing (ST-seq) cohorts.

A. Three PDAC tumor tissues from the GSE233293 cohort, labeled PDAC1, PDAC2, and PDAC3, include HE staining sections, spatial clusters via BayesSpace, count data (nCount), and feature data (nFeature) information.

B. Two PDAC tumor tissues from the GSE202740 cohort, labeled PDAC4 and PDAC5, include HE staining sections, spatial clusters via BayesSpace, count data (nCount), and feature data (nFeature) information.

C. 15 PDAC tumor tissues from the HTAN database, labeled PDAC6 to PDAC20 include HE staining sections, spatial clusters via BayesSpace, count data (nCount), and feature data (nFeature) information.

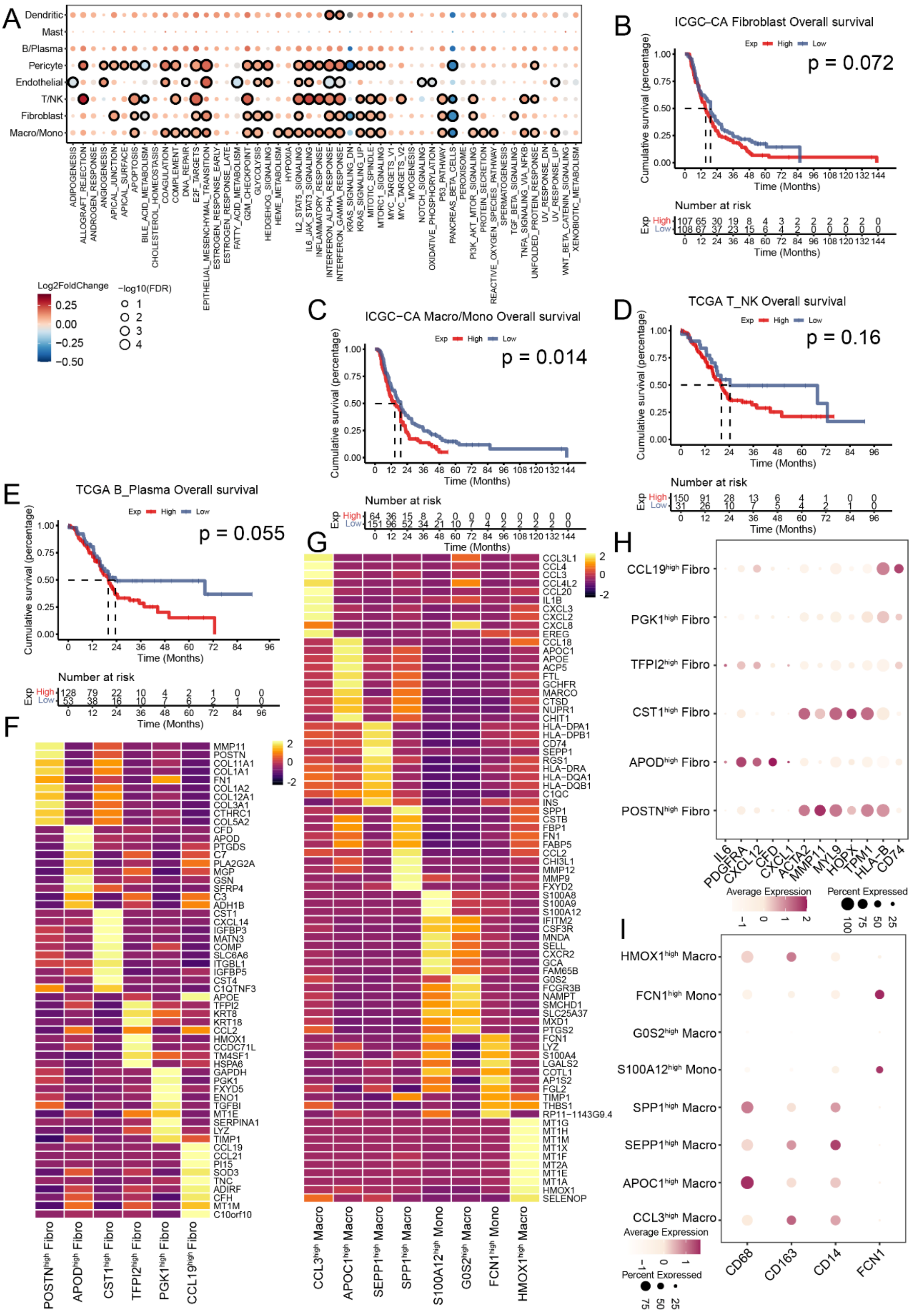

**Figure S3**. Prognosis of PDAC patients and identification of fibroblast and macrophage/monocyte (Macro/Mono) subtypes in the TME.

A. Dot plots showing 50 hallmarks in different cell types between PDAC primary and adjacent/normal tissues. The intensity represents the average fold change of the hallmark GSVA score in tumor versus adjacent/normal tissues. Similar as Figure 2A.The dot size indicates the FDR for each hallmark. Student's t test was used to assess significant differences.

B-E. The Kaplan–Meier curves of fibroblasts in the ICGC-CA cohort (p=0.072; A), Macro/Mono in the ICGC-CA cohort (p=0.014; B), T/NK cells in the TCGA cohort (p=0.16; C), and B/plasma cells in the TCGA cohort (p=0.055; D) within PDAC patient prognosis.

F. The heatmap depicting the expression of representative genes for each of the six fibroblast subtypes.

G. The heatmap depicting the expression of representative genes for each of the eight Macro/Mono subtypes.

H. Bubble plot of the expression of markers for fibroblast types including myofibroblasts, inflammatory fibroblasts, and antigen-presenting fibroblasts across 6 fibroblast subtypes.

I. Bubble plot of the expression of representative markers for macrophages and monocytes across 8 subtypes of Macro/Mono.

A

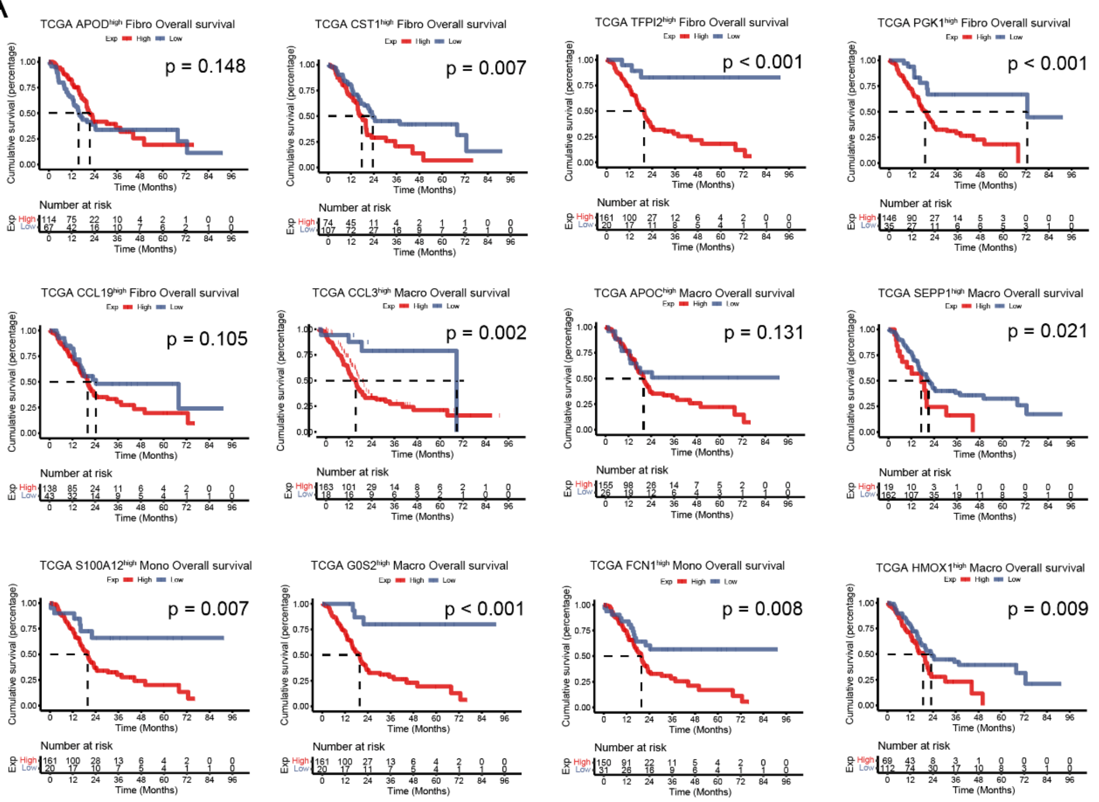


B

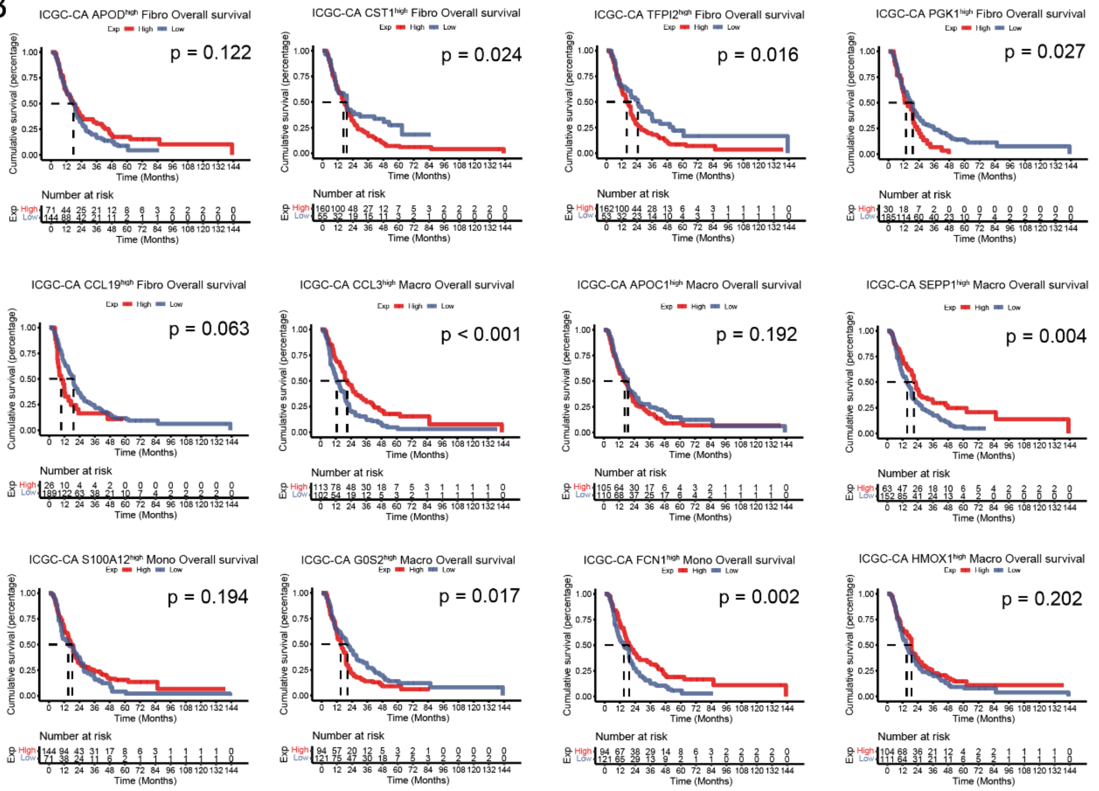


**Figure S4**. Kaplan-Meier curves showing the overall survival (OS) of patients based on subtypes of fibroblasts and Macro/Mono.

A. Kaplan-Meier curves of 5 fibroblast subtypes (excluding POSTN $^{high}$ Fibro) and 7 Macro/Mono subtypes (excluding SPP1 $^{high}$ Macro) with patient OS in the TCGA database.

B. Kaplan-Meier curves of 5 fibroblast subtypes (excluding POSTN $^{high}$ Fibro) and 7

Macro/Mono subtypes (excluding SPP1 $^{high}$ Macro) with patient OS in the ICGC-CA cohort.

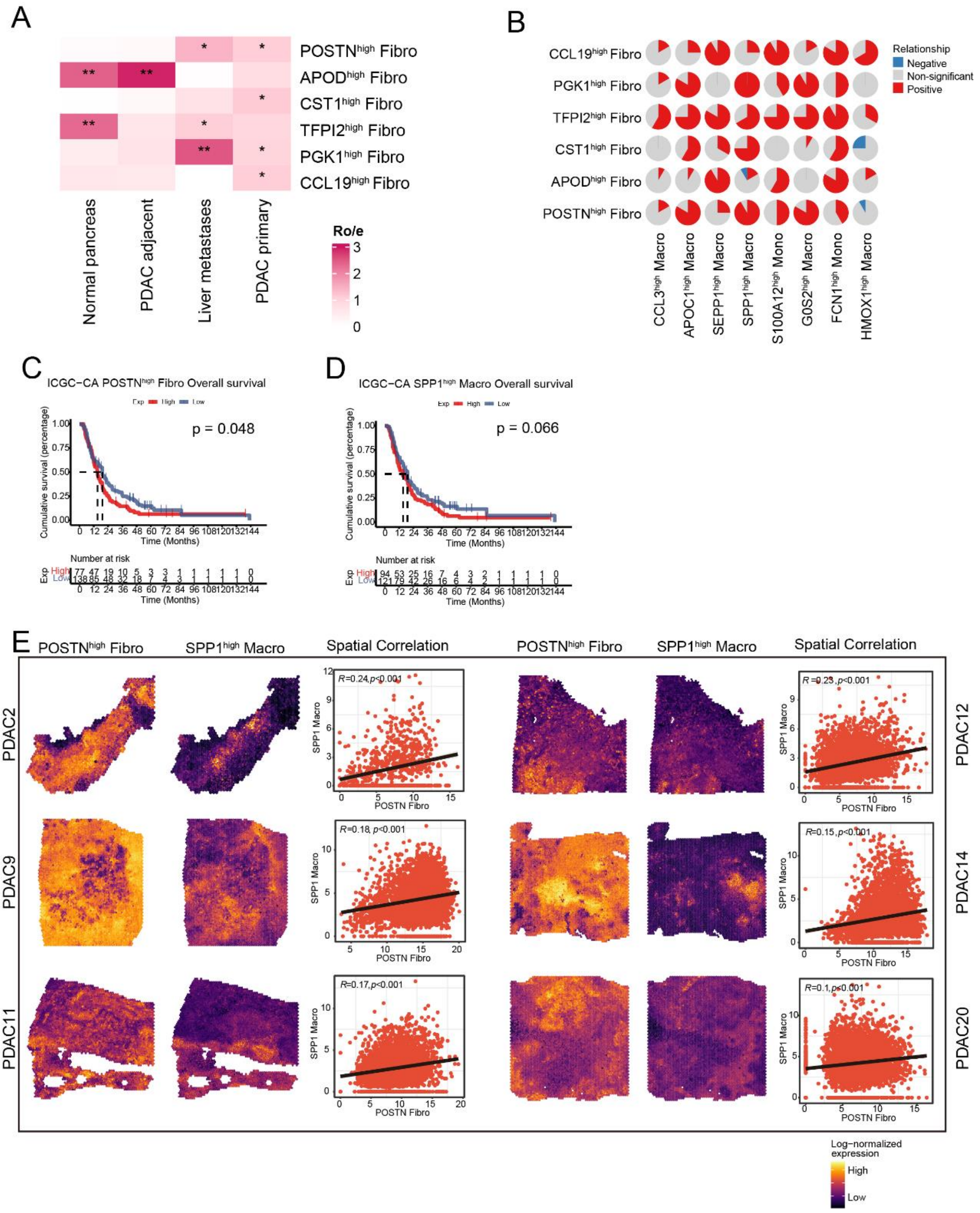


**Figure S5**. Association between POSTN $^{high}$ Fibro subtype and SPP1 $^{high}$ Macro subtype.

A. Heatmap of observed-to-expected ratios (ROE) for different fibroblast subtypes across Sites. P < 0.05 as *, P < 0.01 as **, P < 0.001 as ***.

B. The proportion of PDAC cohorts with positive(red), negative(blue), or non-significant(gray) correlations for the infiltration of pairwise cell types across 12 independent PDAC cohorts.

C-D. Kaplan-Meier curve of infiltration of POSTN $^{high}$ Fibro (p=0.048; B) and SPP1 $^{high}$ Macro (p=0.066; C) cells with patient OS in the ICGC-CA cohort.

E. Expression of POSTN $^{high}$ Fibro subtype and SPP1 $^{high}$ Macro subtype in spatial context and their correlation in ST-seq samples.

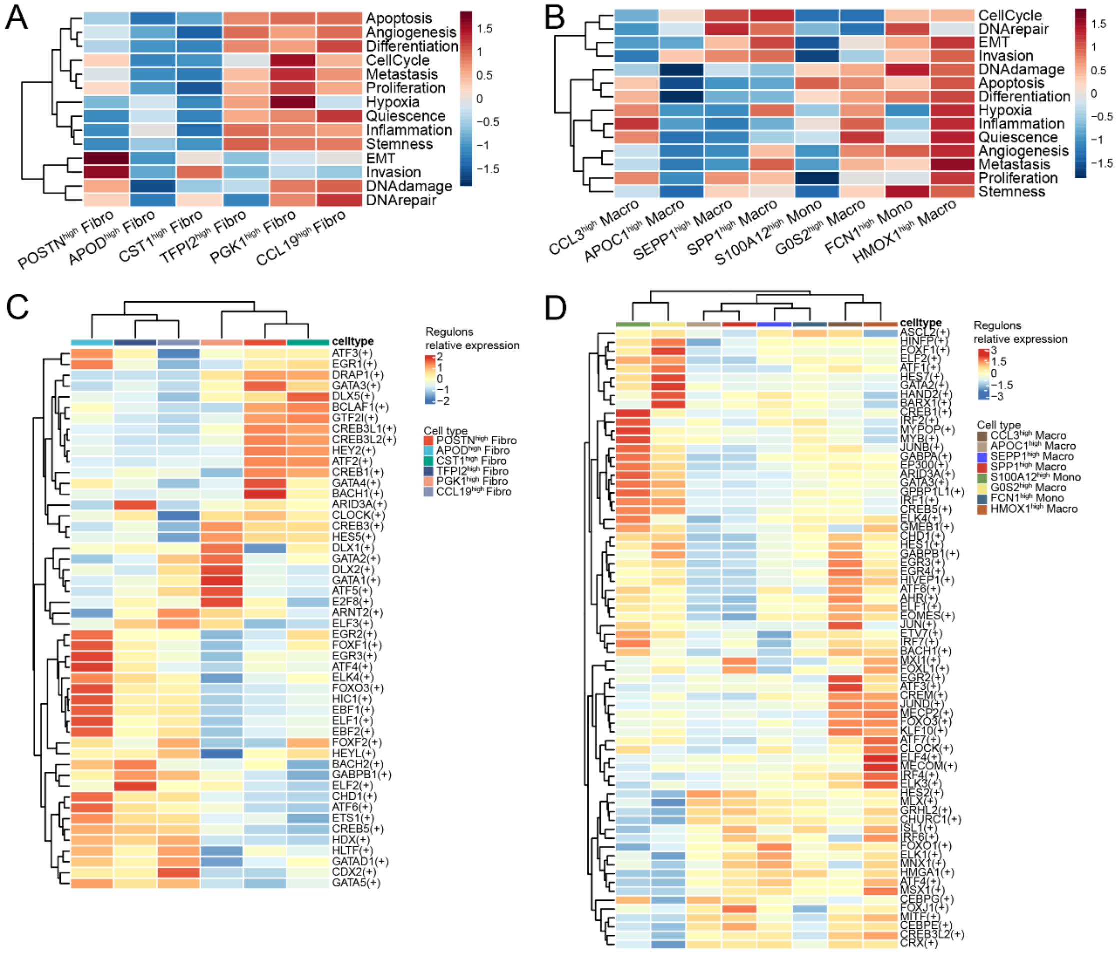


**Figure S6**. Functional enrichment and transcription factor analysis of fibroblast and Macro/Mono subtypes.

A. Heatmap of enrichment scores for 14 cancer-related hallmark pathways across 6 fibroblast subtypes.

B. Heatmap of enrichment scores for 14 cancer-related hallmark pathways across 8 Macro/Mono subtypes.

C. Heatmap of transcriptional regulon activity across 6 fibroblast subtypes.

D. Heatmap of transcriptional regulon activity across 8 Macro/Mono subtypes.

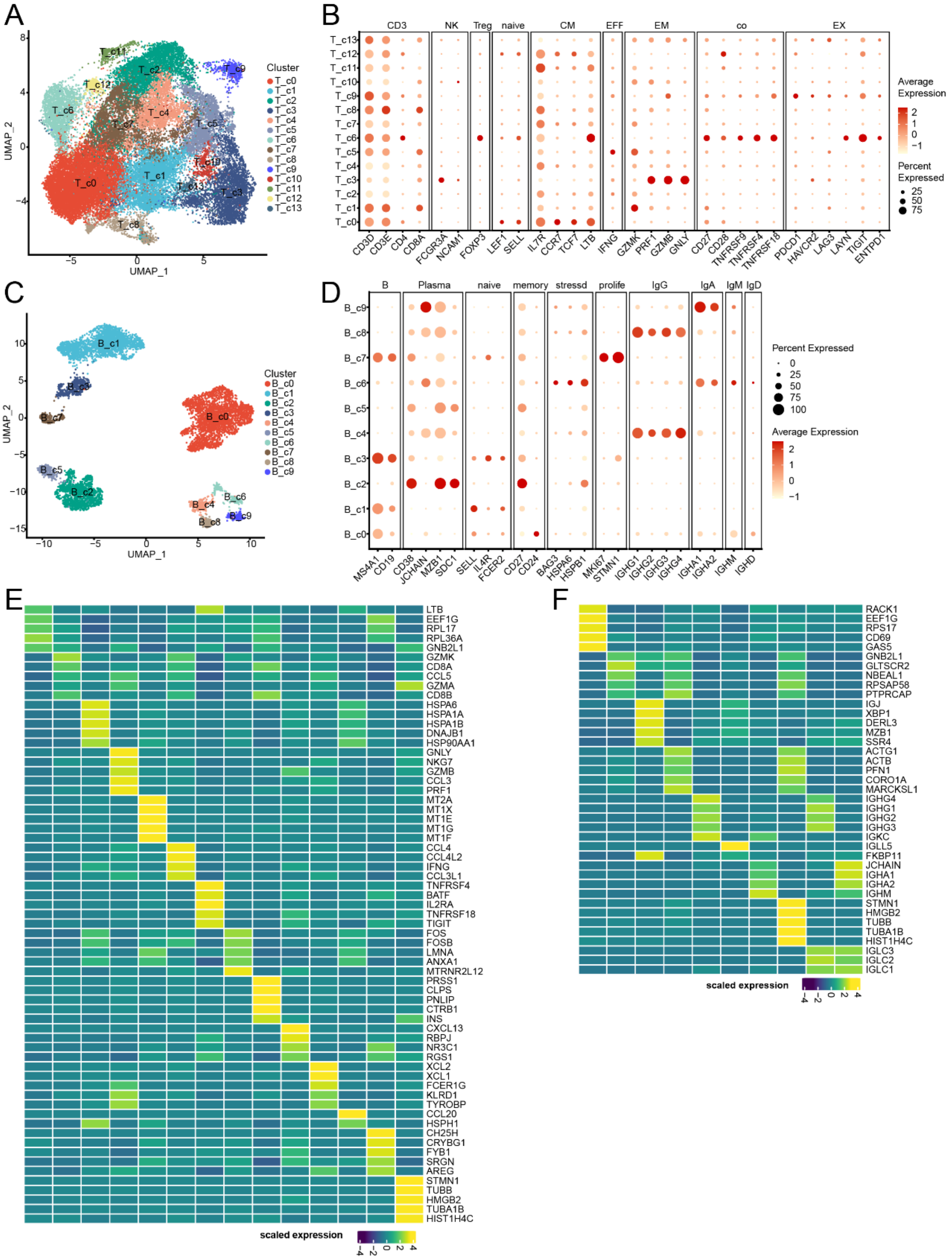


**Figure S7.** Classification and identification of T/NK and B/Plasma subtypes.

A. UMAP plot of the 14 clusters generated by re-dimensioning T/NK cell types.

B. Bubble plot showing the expression of markers CD3, NK, regulatory T cells (Treg), central memory (CM), effector (EFF), effector memory (EM), co-stimulatory (co), and exhausted (EX) across the 14 T/NK clusters.

C. UMAP plot of the 10 clusters generated by re-dimensioning B/Plasma cell types.

D. Bubble plot showing the expression of markers B, Plasma, naive, memory, stressed, proliferative (prolife), IgG, IgA, IgM, and IgD across the 10 B/Plasma clusters.

E. Heatmap showing the expression of representative genes across the 14 T/NK clusters.

F. Heatmap showing the expression of representative genes across the 10 B/Plasma clusters.

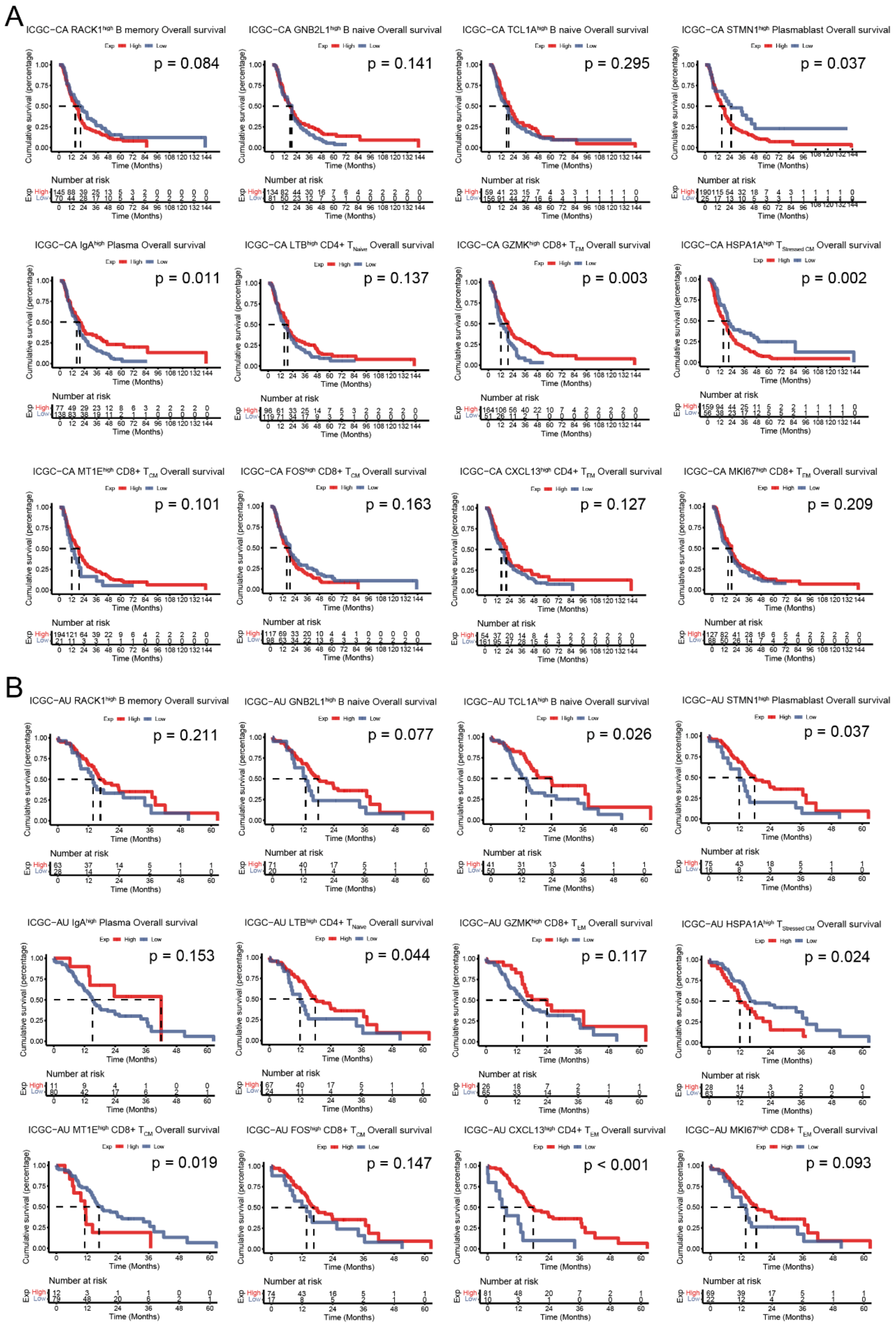

**Figure S8.** Kaplan-Meier curves showing the overall survival (OS) of patients based on subtypes of T/NK and B/Plasma.

A-B. Kaplan-Meier curves of 5 B/Plasma subtypes (including RACK1$_{high}$ B memory, GNV2L1$^{high}$ B naïve, TCL1A$^{high}$ B naïve, STMN1$^{high}$ Plasmablast, and IgA$^{high}$ Plasma cell) and 7 T/NK subtypes (LTB$^{high}$ CD4+ T$_{naive}$, GZMK$^{high}$ CD8+ T$_{EM}$, HSPA1A$^{high}$ T$_{Stressed\ CM}$, MT1E$^{high}$ CD8+ T$_{CM}$, FOS$^{high}$ CD8+ T$_{CM}$, CXCL13$^{high}$ CD4+ T$_{EM}$, and MKI67$_{high}$ CD8+ T$_{EM}$) with patient OS in the ICGC-CA (A) and ICGC-AU (B) cohort.

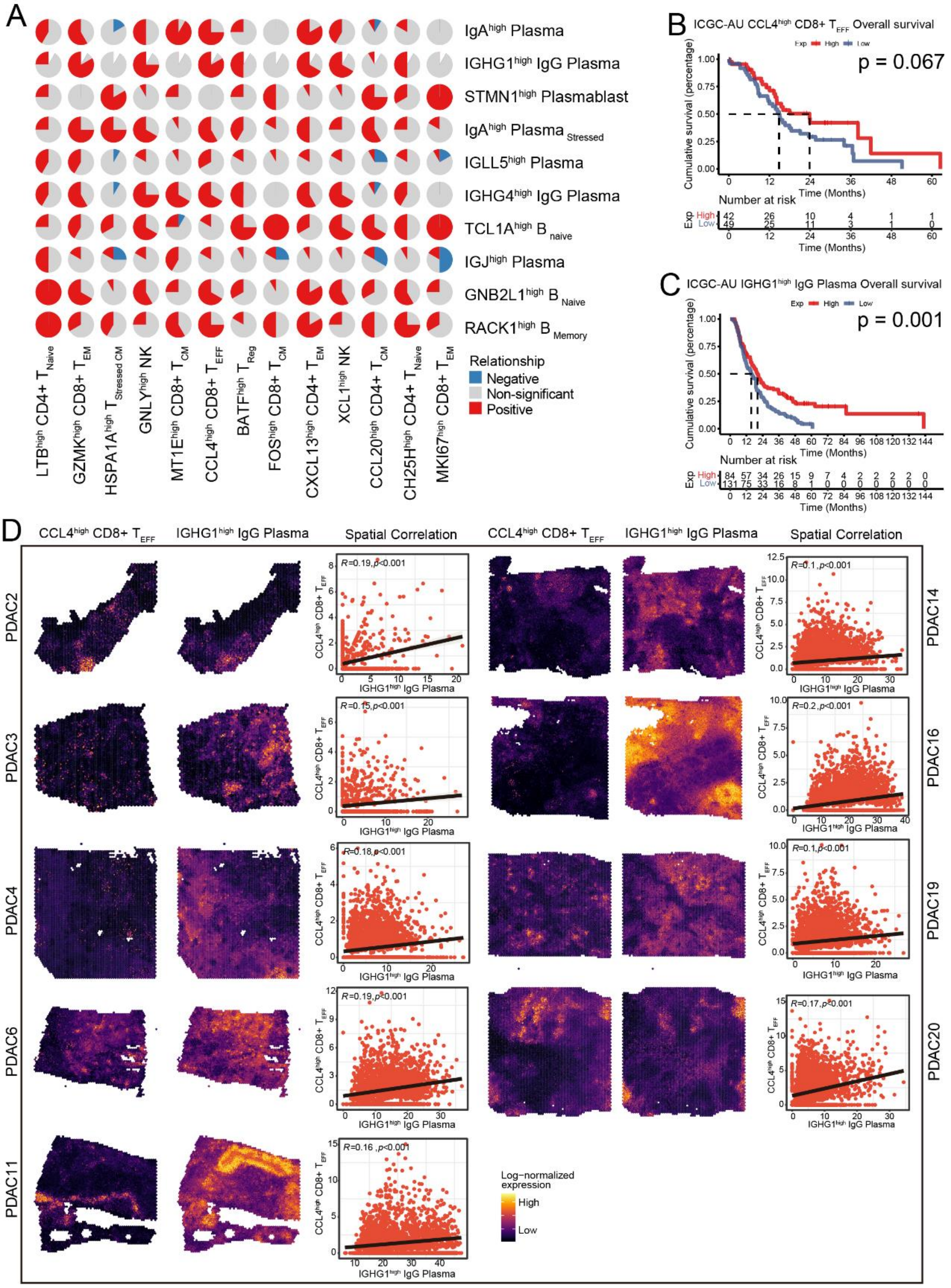

Figure S9. Correlation between CCL4 $^{high}$ CD8+ effector T cells and IGHG1 $^{high}$ IgG plasma cells.

A. The proportion of PDAC cohorts with positive(red), negative(blue), or non-significant(gray) correlations for the infiltration of pairwise cell types across 12 independent PDAC cohorts.

B-C. Kaplan-Meier curve of infiltration of CCL4 $^{high}$ CD8+ T $_{EFF}$ cells (B) and IGHG1 IgG $^{high}$ Plasma cells (C) with patient OS in the ICGC-AU cohort.

D. Expression of CCL4 $^{high}$ CD8+ T $_{EFF}$ cells and IGHG1 IgG $^{high}$ Plasma cells in spatial context and their correlation in ST-seq samples.

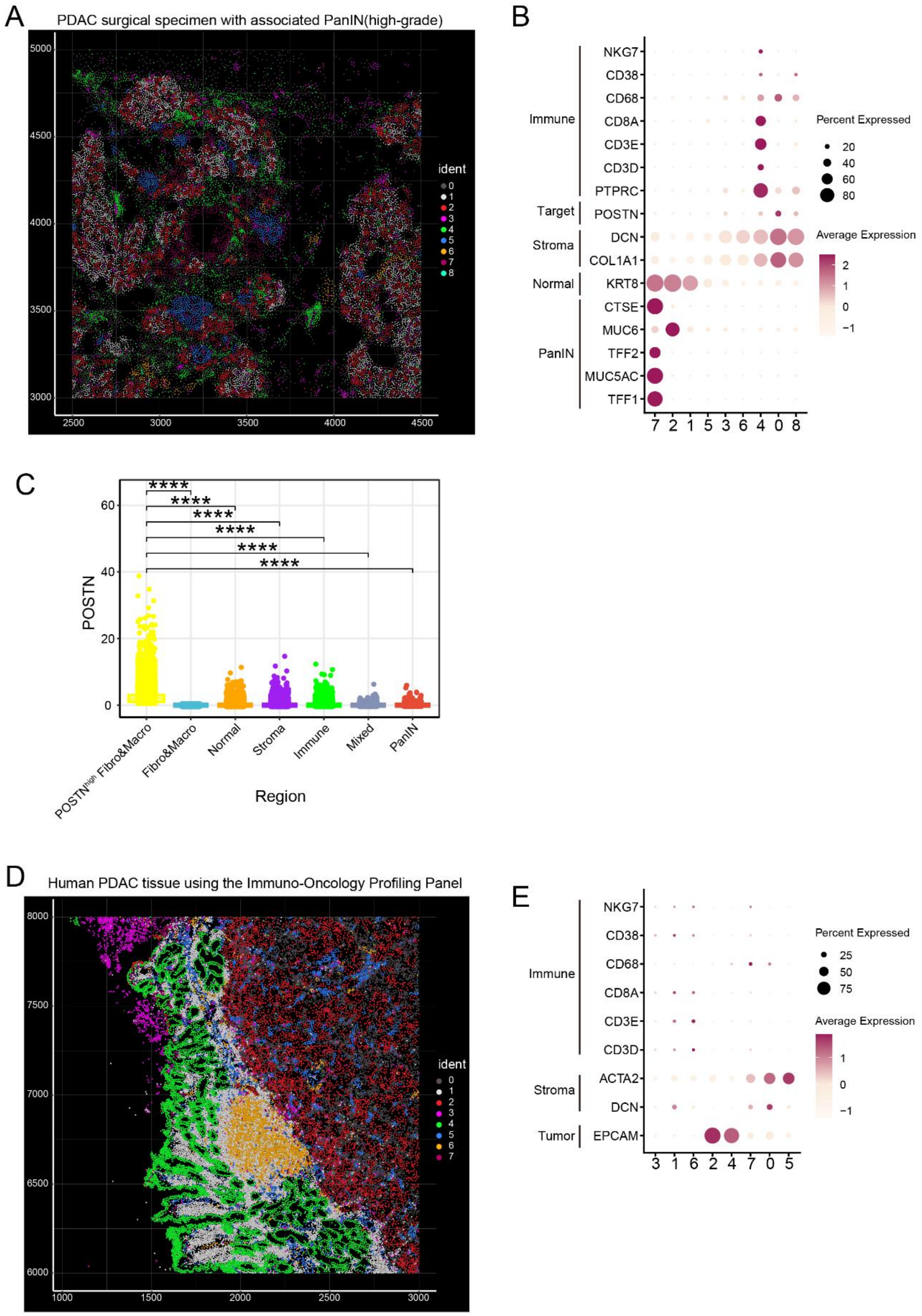


Figure S10. Identification of cellular populations in PDAC tissues using spatial single-cell omics on the Xenium platform

A. Spatial cell cluster distribution map of PDAC surgical specimen with associated PanIN (high-grade) via Seurat V5.

B. Bubble plot showing the expression of selected cell markers for 9 clusters, as shown

in figure A. Markers are categorized into Immune, Target, Stroma, Normal, and PanIN.
C. Scatter plot showing the expression of POSTN across different regions. Wilcoxon test was used to compare the inter-group differences between POSTN high Fibro & Macro region and other regions. P < 0.0001 as ****.
D. An annotated spatial cell cluster distribution map of human PDAC tissue, generated using an Immuno-Oncology Profiling Panel provided by the 10x Genomics dataset.
E. Bubble plot illustrating the expression of selected cell markers for 8 clusters, as shown in figure C. Markers are categorized into Immune, Stroma, and Tumor.

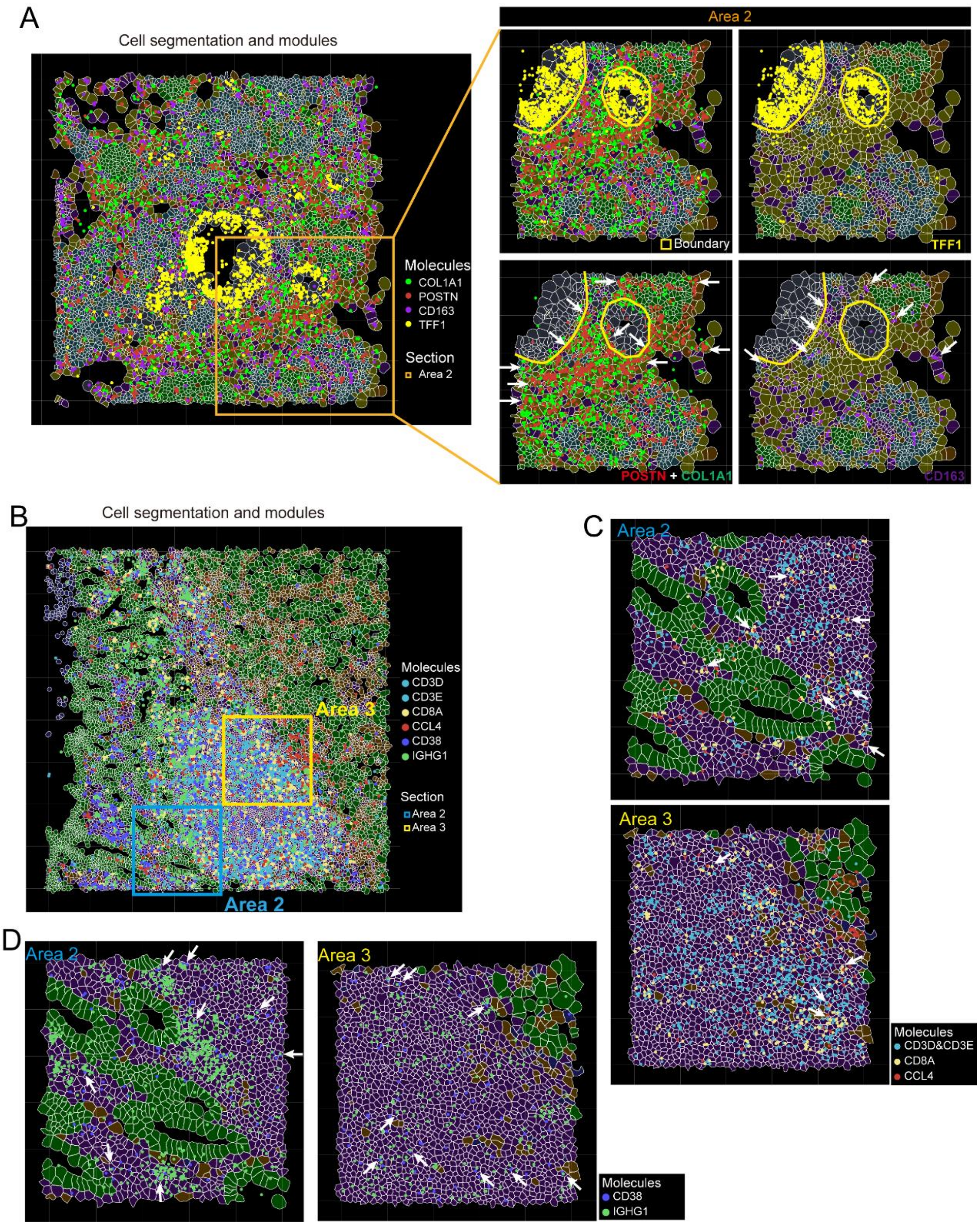


Figure S11. The spatial distribution maps of cell segmentation and modules

A. The spatial distribution maps of cell segmentation and modules in Area2 show Merge, TFF1 (yellow), POSTN+COL1A1(red+green), and CD163(purple), respectively.
B. Spatial distribution of cell segmentation and associated modules in PDAC TME, areas 2 and 3.
C. The spatial distribution maps of cell segmentation and modules in areas 2,3 show CD3D&CD3E (blue), CD8A (yellow), and CCL4 (red), respectively.
D. A spatial distribution maps of cell segmentation and modules in areas 2,3 show CD38 (purple), and IGHG1 (green), respectively.
